# A deep learning approach to capture the essence of *Candida albicans* morphologies

**DOI:** 10.1101/2021.06.10.445299

**Authors:** V Bettauer, ACBP Costa, RP Omran, S Massahi, E Kirbizakis, S Simpson, V Dumeaux, C Law, M Whiteway, MT Hallett

## Abstract

We present deep learning-based approaches for exploring the complex array of morphologies exhibited by the opportunistic human pathogen *C. albicans*. Our system entitled Candescence automatically detects *C. albicans* cells from Differential Image Contrast microscopy, and labels each detected cell with one of nine vegetative, mating-competent or filamentous morphologies. The software is based upon a fully convolutional one-stage object detector and exploits a novel cumulative curriculum-based learning strategy that stratifies our images by difficulty from simple vegetative forms to more complex filamentous architectures. Candescence achieves very good performance on this difficult learning set which has substantial intermixing between the predicted classes. To capture the essence of each *C. albicans* morphology, we develop models using generative adversarial networks and identify subcomponents of the latent space which control technical variables, developmental trajectories or morphological switches. We envision Candescence as a community meeting point for quantitative explorations of *C. albicans* morphology.

## Introduction

Fungal infections represent an urgent and significant threat to human health affecting 1.2 billion people yearly^1–3^. They kill approximately the same number of people (1.6 million) as malaria^4^, and are implicated in cancer progression^5^. *Candida albicans* is one of the most important of these human pathogens^6^ and carries a significant socio-economic burden^2,7–9^. As such, it has become an important system for studying fungal pathogenicity. *C. albicans* is morphologically classified as a pleomorphic yeast-like fungus and systematically classified as an ascomycete. It is well adapted to its role as an opportunistic pathogen with a diverse range of morphologies that predominate in different niches. The morphologies can be broadly partitioned into vegetative and mating-competent forms^10,11^. We provide a brief review of the morphologies central to our effort here (**Figure 1A-E**).

**Figure 1.**
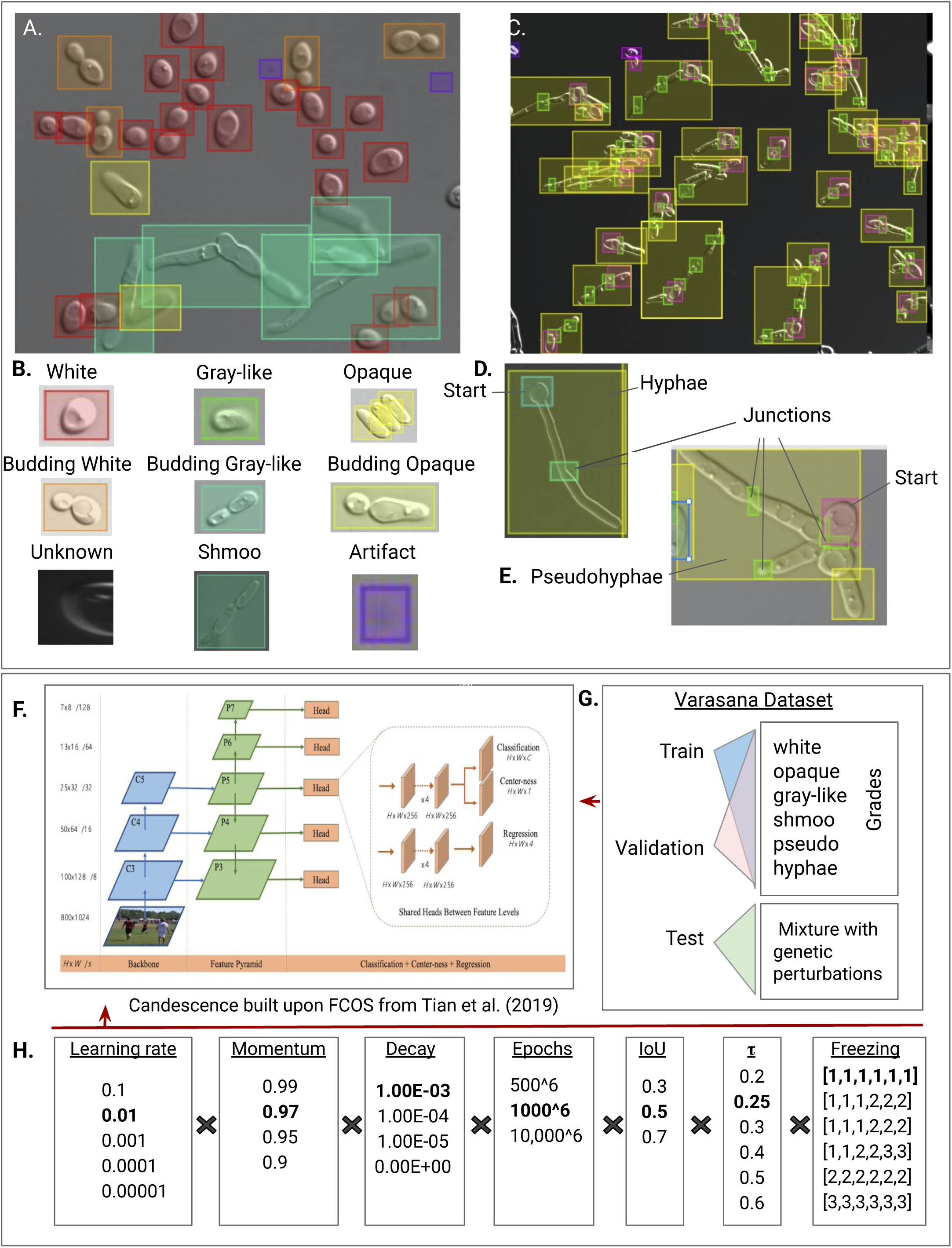
**A.** An example of a typical DIC image in Labelbox, a learning-set creation tool for group annotation. The color codes for bounding boxes described in panel **B.** The artifact class is used to bound imperfections and technical artifacts in the images, whereas the unknown class is used to bound cells for which we could not judge morphology. **C.** Labelling of an image that is enriched for pseudohyphae. Note that the overall intensity of our images are not required to be the same. **D.** Given the size and complexity of the filamentous forms, we annotated each hyphae or pseudohyphae using three classes. **D, E.** H-start and P-start are intended to label only the “start” of the hyphae or pseudohyphae. Since junctions are different between hyphae and pseudohyphae, we labelled them with the H- and P-junction classes. Finally we bounded the entire filamentous object with a rectangle; note that such bounding boxes overlap with surrounding cells. **F.** Our system is built upon the FCOS software with structure unaltered from Tian et al. (2019). **G.** The Varasana learning set is first partitioned into three components (training, validation, and testing). Then both validation and testing are split into six grades. The grades are ordered from white to hyphae. When an image appears first in grade i, it is also included in all grades j > i. The test set contains examples of both wildtype SC5314 and SN148a cells, but also a collection of genetic and environmental perturbations that induced abnormal *C. albicans* morphologies. **H.** Using the Varasana learning set, we performed a grid search across several hyperparameters (n=∼30K combinations). Bold face indicates the combination of hyperparameters that had the best performance.

<u>Vegetative forms</u> differ in cell and colony morphologies, their molecular state and their behaviors as infectious agents. These yeast forms are commonly found on mucosal and skin surfaces where they grow benignly and are tolerated by the host immune system^12^. The *white form* has a round unicellular morphology (**Figure 1A,B**). If engulfed by macrophages, the white yeast will switch to the hyphae morphology as a means of immune escape^13^. The *opaque form* is larger and has a more rectangular shape than white cells (**Figure 1A,B**). *C. albicans* can mate only when both cells are in the opaque morphology^14^. Switching between the two morphologies is rare (∼10^4^ cell divisions), stochastic, and strongly influenced by environmental cues and homozygosity of mating type. White cells are believed to be better suited to internal infections, while opaque cells thrive in skin infections^15^. The *gray morphology* is an alternative, stable form between the white and opaque morphologies (**Figure 1A,B**). Gray cells are “shiny” and small like white cells yet elongated like opaque cells in solid media^16^.

*C. albicans* can also assume <u>mating-competent</u> morphologies including the budding yeast white cell (**Figure 1B)** and two distinct <u>filamentous</u> forms (**Figure 1C-E**). The *hyphal form* is characterized by long tube-like filaments without constrictions at the site of septation. Hyphae are able to invade epithelial and endothelial cells, and damage host tissue in mucosal infections in order to gain access to the bloodstream^17^. The yeast to hyphal switch is initiated under a variety of environmental conditions including presence of serum, neutral pH, 5% CO2, N-acetyl-D-glucosamine, amongst others^18^. In the first cell cycle, the germ tube morphology is observed, manifesting as tube projecting from the round yeast cell (**Figure 1D**). The second filamentous growth form, *pseudohyphae,* are significantly different from hyphae at the molecular level in terms of cellular regulation and growth^19–21^ but also in terms of their phenotypic presentation. Pseudohyphae tend to have more branch points, because mother-daughter attachments are more easily disrupted than in hyphae (**Figure 1E**). Unlike hyphae, pseudohyphal cells exhibit synchronous cell divisions and septation at the mother-bud neck^22^. Weak filament-inducing conditions (e.g. high-phosphate medium) favour a pseudohyphae over a hyphae morphology. Like hyphal cells, they interact with the mouth, vagina and bloodstream of the host. The *shmoo morphology* arises from opaque cells which have formed a projection produced by the a and ɑ types when preparing to mate, leading to tetraploid zygote formation.

Some less frequent *C. albicans* morphologies are not considered in our effort here. This includes *chlamydospores* which are formed at the end of pseudohyphae and hyphae filaments (suspensor cells). They have a thicker cell wall and are larger than blastospores. The final chlamydospore has an elaborate septin-derived substructure. We also do not consider *Gastrointestinally-IndUced Transition* (GUT) cells, which are derived from the white morphology when passaging through the gut of the host. The morphology confers survival in the digestive tract via metabolic adaptations to available nutrients in this environment^23^. The *trimera* morphology is sometimes observed after unequal chromosome segregation under, for example, stress from antifungal exposure. Finally, we do not consider the *goliath* morphology which arises in response to mammalian host nutritional immunity strategies including zinc sequestration^24^. When *C. albicans* is unable to scavenge host zinc during endothelial invasion, the fungus transforms to a giant yeast cell morphology with advanced adhesion, a property shared with the hyphal morphology.

There have been several earlier efforts in fungal image analysis^25^ including approaches to computationally differentiate between types of fungi and allergenic fungal spores^26^, and to characterize the macro-structure of mycelium or filamentous growth^27–34^. Tleis and Verbeek experimented with a suite of machine learning techniques to segment *S. cerevisiae* cells and measured a range of features and textures from two channel images acquired by a laser scanning confocal microscope^35^. Wang et al. provided a segmentation method with an efficient edge-tracing algorithm for bright-field images of fission yeast, which have a consistent oblong morphology^36^. Another effort employed microfluidics to capture individual *S. cerevisiae* cells; non-fluorescent images were used to train a classifier of cell cycle state for each cell^37^.

Deep learning was first applied to biomedical imaging in 2012 with the approach from Ciresan and colleagues^38^ to automatically segment neuronal structures depicted in stacks of electron microscopy images. Shortly thereafter, Rosenberger et al.^39^ developed a generic convolution network (termed U-net) for image segmentation that has since been used in many biomedical-related image recognition challenges. Our work here builds upon a different but well-established deep learning architecture entitled Res-Net^40^. These and other deep learning-based approaches have been extended to cell counting, detection and morphology. To date, these methods have been primarily used in the context of human tissue analyses^41–44^.

Deep learning has opened up many new avenues of investigation in microscopy through the introduction of new techniques for image transformation, object localization in super resolution computing, and cross-modality imaging^45^. One challenge is to establish a model of the morphology of a single cell, and then detect when cellular perturbagens change that morphology^46–48^. Approaches use both generative adversarial training^49^ and variational autoencoder-based methods^50^. Recently, a very powerful form of deep learning-based image analysis entitled Faster Region-based Convolutional Neural Network (Faster R-CNN)^51^ was applied to images of blood smears from individuals infected with malaria^52^. The goal was to identify different human cell types concomitantly with the protist *Plasmodium vivax* in images.

With respect to fungi, there have been significant advances to integrate robotic-controlled hardware with high-content image analysis tools to rapidly screen and analyze hundreds of thousands of images of *S. cerevisiae* to, for example, identify essential genes and genetic interactions^53,54^. These works were the first to bring large-scale image analysis at the subcellular level; the first deep learning approach (DeepLoc) expanded on this subcellular microscopy work in *S. cerevisiae*^55^.

Our effort investigates two novel inter-related image recognition challenges with *C. albicans* using methodology from deep learning. We first develop a system that can automatically detect *C. albicans* cells from microscopy images and label each detected object with its morphology. The model is trained using our large compendium of Differential Interference Contrast (DIC) images containing individual *C. albicans* cells in different morphological states. We treat this problem as a multi-class, multi-object object detection problem where 15 distinct classes are used to describe nine morphologies. Some images contain as many as one hundred tightly packed cells. Our method builds upon a fully convolutional one-stage object detector (FCOS)^56^ and a novel cumulative curriculum-based learning strategy that stratifies images by difficulty. Then, building upon our ability to accurately identify and classify cells, we develop a deep learning-based model that captures the essence of each *C. albicans* morphology. These models, which are based upon generative adversarial networks (GANs), can be interrogated to identify components of its latent space that control various features of the images. This includes technical variables but also biologically-relevant processes such as developmental trajectories or transitions between morphologies. These models provide the first dynamic and continuous approach to capturing the essence of the different morphological states and transitions between them. We show how these models can be used to automatically identify subtle changes from wild-type phenotype when given images from genetically perturbed *C. albicans* populations.

## Results

### A deep learning approach to recognizing *C. albicans* morphologies

Our first goal is to develop a fully automated tool that identifies the location and morphology of cells from microscopy images of *C. albicans* populations. We cast this challenge as a multi-object, multi-class detection problem: (i) the system identifies the location of all *C. albicans* cells in each image, and then (ii) predicts the morphology of each hypothesized cell. Images may contain an arbitrary number of individual objects (cells or artifacts). Our system classifies nine of the 12 reported *C. albicans* morphologies: yeast white, budding yeast, opaque, budding opaque, shmoo, gray-like, budding gray-like, hyphae, and pseudohyphae. Each detected object is assigned the morphology with the highest likelihood, provided the probability of the prediction exceeds a user-defined threshold τ.

Our software (entitled *Candescence*) is based on what is termed a fully-convolutional one-stage object detector (FCOS)^56^, a new approach that has several benefits over other deep learning algorithms for multi-object, multi-class problems (**Figure 1F**). One of its key advantages resides from how it flags areas of an image likely to contain an object. Rather than requiring many parameters, which collectively control the size, location and total number of bounding boxes, the FCOS considers each individual pixel in an image as potentially centering an object, abating the need to optimize many hyperparameters simultaneously. FCOSs are able to handle objects of variable size, an important property given the difference between, for example, yeast white and hyphal cells.

FCOSs can make use of transfer learning, a technique where a neural network is first trained in a distinct but similar context compared to the problem at hand. In our case, we begin with a FCOS trained with the ResNet-101 dataset, a convolutional neural network trained on more than a million images partitioned into 1000 classes of common household objects and animals^40^. Intuitively, transfer learning guarantees that our neural network comes equipped with the basic circuitry to recognize simple shapes, shades, edges and textures. Candescence allows this set of rich feature representations to be retrained for our morphologies, requiring only a tractable number of *C. albicans* images. **Methods 1** describes the architecture and technical parameters of the FCOS.

### An image compendium of *C. albicans* morphologies

Supervised machine learning problems including this image recognition problem exploit a learning set, which is a curated collection of images where bounding boxes have been drawn and labelled for every object. We constructed such a learning set by growing *C. albicans* SC5314 or SN148A colonies under conditions necessary to induce each of the nine morphologies (**Table 1A, Methods 2**): yeast white, budding white, opaque, budding opaque, gray-like, budding gray-like, shmoo, hyphae, and pseudohyphae. The number of observed goliath, zygote, chlamydospores and trimera cells was too small for computational training and were excluded from the analysis. This first version of our software also did not consider *C. albican*s GUT cells as they require special culturing conditions. Each population was stained with calcofluor white and prepared for DIC and fluorescent microscopy at various magnifications (40x-100x; **Methods 3**). We also cultured and imaged *C. albicans* colonies that had specific genetic perturbations known to affect morphology. In total, 1214 images were generated (**Supplemental Table 1**).

**Table 1.**
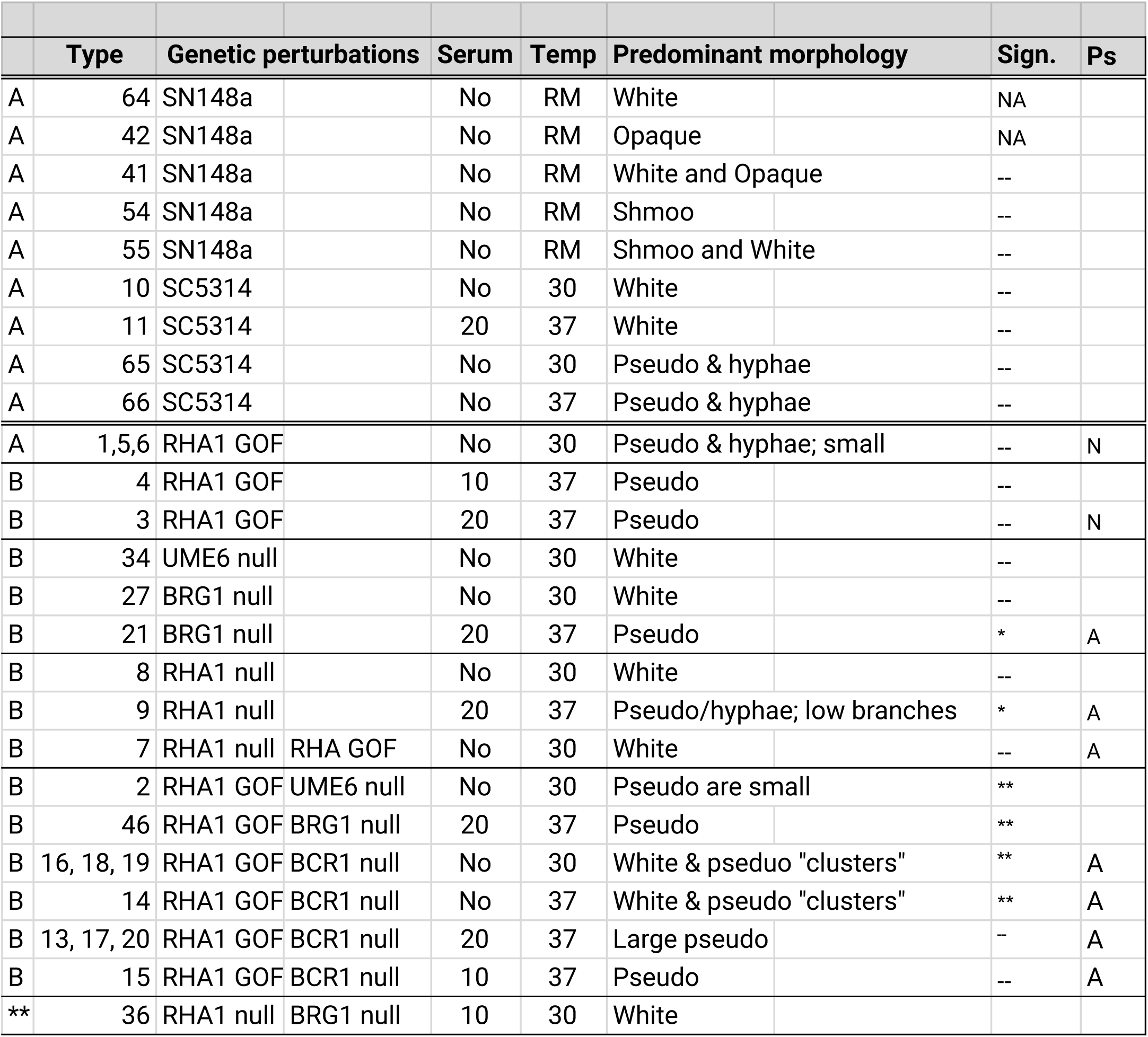
Summary of the Varasana learning set from Supplemental Table 1. An A indicates that some images from this type of colony were used in the training and validation datasets, and B indicates that some images were used in the test set. All genetic perturbations were performed in SC5314 cells. In the Serum column, a No indicates only YPD media was used, otherwise the percent serum added to YPD is recorded. RM indicates room temperature. The significance column indicates the results of applying the proportionality test to determine if there was a difference in performance between the validation and test set. Here * and ** indicate p-values below 0.05 and 0.01 respectively and -- indicate a p-values > 0.05. In cases where a p-value is provided for a type A colony, only images that were omitted from the training and validation datasets were used.

In each of the resultant DIC images, a bounding box was manually drawn around each *C. albicans* cell and labelled with its morphological class following the guidelines of Sudbery et al.^21^, Whiteway and Bachewich^11^, Tao et al.^16^ and Noble et al.^23^ (**Methods 4**). More specifically, cells with a round-to-oval morphology (4.9μm × 6.8μm) were labelled *white* unless they were attached to a second smaller white cell with concomitant evidence of a bud site (as sometime assisted by manual inspection of the calcofluor white signal in the matched fluorescent image); in this case they were labelled *budding white* (**Figure 1A,B**). Ellipsoidal cells approximately twice the size of white cells (9.5μm × 11.8μm) were labelled *opaque*. Like white cells, they were labelled *budding opaque* if there was evidence of a smaller opaque cell with a new bud site. Cells were labelled *shmoo* if they had an irregular (often boomerang) shape and large vacuoles. Many of our images contain a significant number of vegetative-like cells that are visibly distinct from both white and opaque morphologies. These cells are ellipsoidal in shape, with a size in the upper quantile of white cells, but below the size of opaque cells. These properties are close in spirit to the *gray* morphology of Tao and colleagues^16^, although we did not grow *C. albicans* colonies under conditions that specifically induce this morphology. To date, confirmation of the gray cell morphology requires the use of specific molecular markers. However, given the distinctiveness and ubiquitousness of these cells in our data, we decided to explicitly model this structure and use the term *gray-like*.

The diversity and complexity of filamentous cells necessitated the development of an approach that uses several markers concomitantly to reliably identify them. Hyphae are thin tube-shaped cells with a width of ∼2μm. Pseudohyphae are multicellular entities which tend to be elongated and ellipsoidal in structure. The minimum width is 2.8μm. Our markup scheme first bounds the entire hyphae or pseudohyphae. Since filamentous cells are often long and irregular in shape, the bounding box almost always overlaps, or completely subsumes, the bounding box of other objects. Since this overlap represents a significant challenge for learning algorithms, we also labelled the location of the original germ bud of each hyphae, or estimated the start cell of the pseudohyphae; these bounding boxes were labelled *H-* and *P-start* respectively. Compared to hyphae/pseudohyphae bounding boxes, H- and P-start are much smaller, and therefore disjoint from other objects in the image (**Figure 1C-E**). As is the case for the start sites, the septal junctions in hyphae and pseudohyphae are visually distinct from one another. Only pseudohyphae have constrictions at the mother-bud neck and subsequent septal junctions. We placed bounding boxes at these junctions and labelled them as *H-* or *P-junction*. Those cells for which we could not reach agreement on morphology were labelled as *unknown*; non-cellular events in the images were labelled as *artifacts*.

We stress here that our strategy attempted to label each cell using only its visual appearance and independent of other factors including the predominant morphology of cells in the image. For example, in an image of a colony grown under conditions which will enrich for yeast white cells, we still expect one in 10^4^ cells to stochastically assume the opaque morphology; cells on the border between these morphologies may have been biased towards the background yeast white form. In some cases, a consensus was difficult to achieve across labellers.

### Varasana: a cumulative curriculum-based *C. albicans* learning set for Candescence

Learning sets are typically tri-partitioned (**Figure 1G**). The *training set* contains a large set of images which are presented to the neural network during training and used to fit the model (i.e. update the weights of each arc in the neural network). The *validation set* is typically a smaller sample of data which is used to generate an unbiased estimation of the model fit during training. It provides a means to tune the hyper-parameters of the model. The *test dataset* is used only once after all training is complete to assess the final fit of the model. We partitioned the images from **Table 1A** so that cells from each morphology are assigned to the training and validation set at a 7:3 ratio. This procedure was confounded by the fact that there is significant variability in both the number of cells per image and the composition of morphologies per image. From the collection of 1214 images, the training and validation dataset contains 216 and 94 images respectively. These images in turn contain 4880 and 1958 objects respectively. The independent test contains the remaining 904 images.

FCOSs have several hyper-parameters that affect the overall performance of the system including the learning rate (the amount weights are updated during gradient descent), momentum (a parameter that stipulates how many previous steps can be used to determine the direction of a weight update), decay (a regularization parameter restraining the complexity of our model), epoch number (the number of times the learning phase cycles through the complete training and validation sets), the IoU (the intersection of union statistic controls how accurate the regression of the bounding box must be), the threshold τ (the lower bound for the probability an object is assigned a class), and others (**Figure 1H**). We performed a grid search across a range of values for these parameters and measured convergence, model complexity and performance after each trial (**Methods 4-5**). Although the resultant classifiers had good performance for some morphologies (e.g, F1 ∼78% for white, budding white, opaque, gray and shmoo), several classes remained poorly predicted (e.g. F1 ∼50% for pseudohyphae and hyphae related classes) and budding classes were poorly distinguished from their parent (e.g. F1 ∼60% for budding white, gray and opaque).

The steep increase in difficulty between yeast white and the filamentous classes highlighted the need for a more structured learning set. We opted for a curriculum approach^57,58^, a well-established concept in psychology which has re-emerged in the deep learning community. The fundamental idea is to structure the learning set so that the neural network is exposed to concepts according to their difficulty, with the easiest concepts presented first. Using the results from our original grid search to judge easy and difficult objects, we re-designed our learning set into grades one through six. Each grade was enriched for a specific subset of classes, although at least one example of all classes was present at each grade. Simple examples of yeast white for example were placed in grade 1, and more complicated examples (e.g. crowded images, poor image quality) were placed in later grades. In general, grade 1 is enriched for yeast white and budding white cells, grade 2 introduces opaque and budding opaque, grade 3 presents gray and budding gray, grade 4 focuses on the shmoo form, grade 5 on pseudohyphae and grade 6 concludes with hyphae. Once an image appears at a certain grade, it appears in all subsequent grades. We hypothesize that this cumulative strategy ensures that lessons learnt early with populous morphologies such as yeast white and opaque are retained when the complicated filamentous morphologies are presented to the learner. All grades had many examples of artifacts and unknowns. To the best of our knowledge this is the first use of a cumulative curriculum learning approach.

A new grid search was conducted but this time the hyper-parameters were allowed to vary across the different grades. This grid search also considered different levels of freezing in our four layer FCOS. Freezing refers to the process of disallowing layers of the neural network to change. For example, the most restrictive freezing regimen disallows any of the four layers to change in response to new examples, implying that our classification is solely based on the original transferred ResNet-101. The most permissive freezing strategy allows all layers to be updated during training when presented with our images. **Supplemental Figure 1** breaks down the learning set by grade and classes, and provides the distribution of the number of cells per image.

### Searching for classifiers of *C. albicans* morphologies

A second grid search was performed with the validation dataset across the hyperparameter space. The search required ∼60 days of continuous computation on a system with 10 GPUs. We judged performance initially using the mean average precision (mAP)^59^, a standard approach for measuring the performance of multi-object/multi-class problems, across all trials with a final value of 0.407 (**Methods 5**). This best classifier has a learning rate of 0.01, momentum of 0.97, and decay of 0.001 (**Figure 1H**). Analysis of the loss curves established that 1,000 epochs at each grade sufficed for convergence for each of the three component loss curves (**Methods 5, Supplemental Figure 2**). An initial performance assessment suggested that an IoU of 0.5 and a τ of 0.25 maximized the recall, precision and F1 which were estimated to be 82.4%, 66.5% and 73.7% respectively (**Table 2**).

**Table 2.**
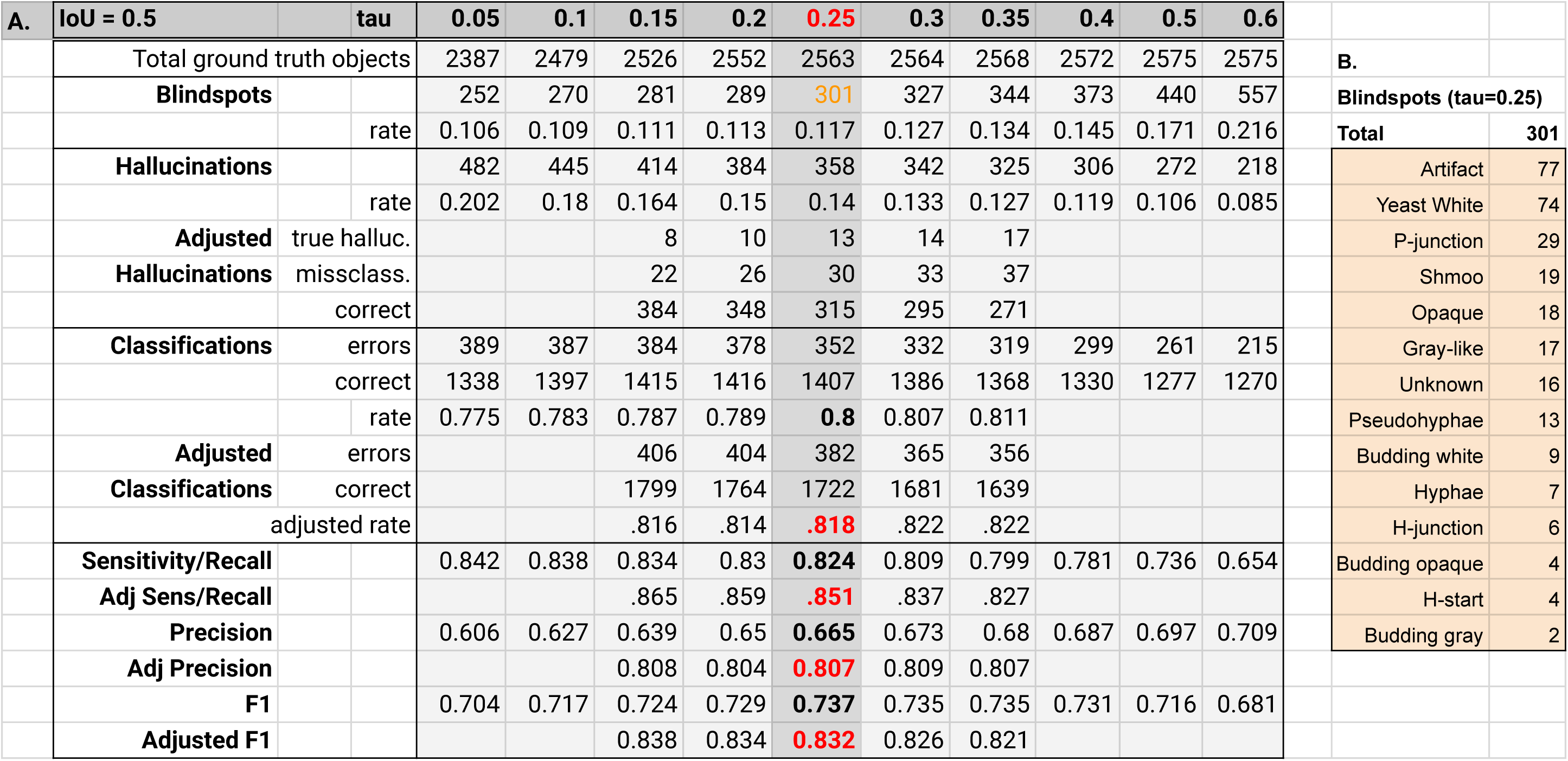
Exploration of the performance of the FCOS in the grid search. Panel **A** provides the relevant summary statistics for blindspots, hallucinations and classifications. With *a posterior* analysis and correction of the hallucinations, performance was recalculated and reported in the Adj Sens (adjusted sensitivity) column. True positives correspond to a correctly identified object that is also correctly classified. In an FCOS, all pixels that are not part of a bounding box and which are not predicted to be part of a bounding box are true negatives. False positives are hallucinations and incorrect classifications. False negatives are blindspots only. **Panel B** reports the blindspots per class for the best performing classifier at τ=0.25.

Somewhat surprisingly, the performance of classifiers with more liberal freezing strategies was better than strategies that froze layers. We hypothesize that the cumulative nature of our learning set guaranteed that the network retained learnt rules from early grades without the need for freezing. To the best of our knowledge, we have not seen any exploration of cumulative curriculum learning nor the observation that a cumulative curriculum learning approach may suffice in lieu of freezing. **Figure 2** depicts the ground truth labels (left) and predicted objects and their classifications (right) across three typical images containing many of the morphologies. The images help to show that Candescence is able to cope with overlapping objects in dense images.

**Figure 2.**
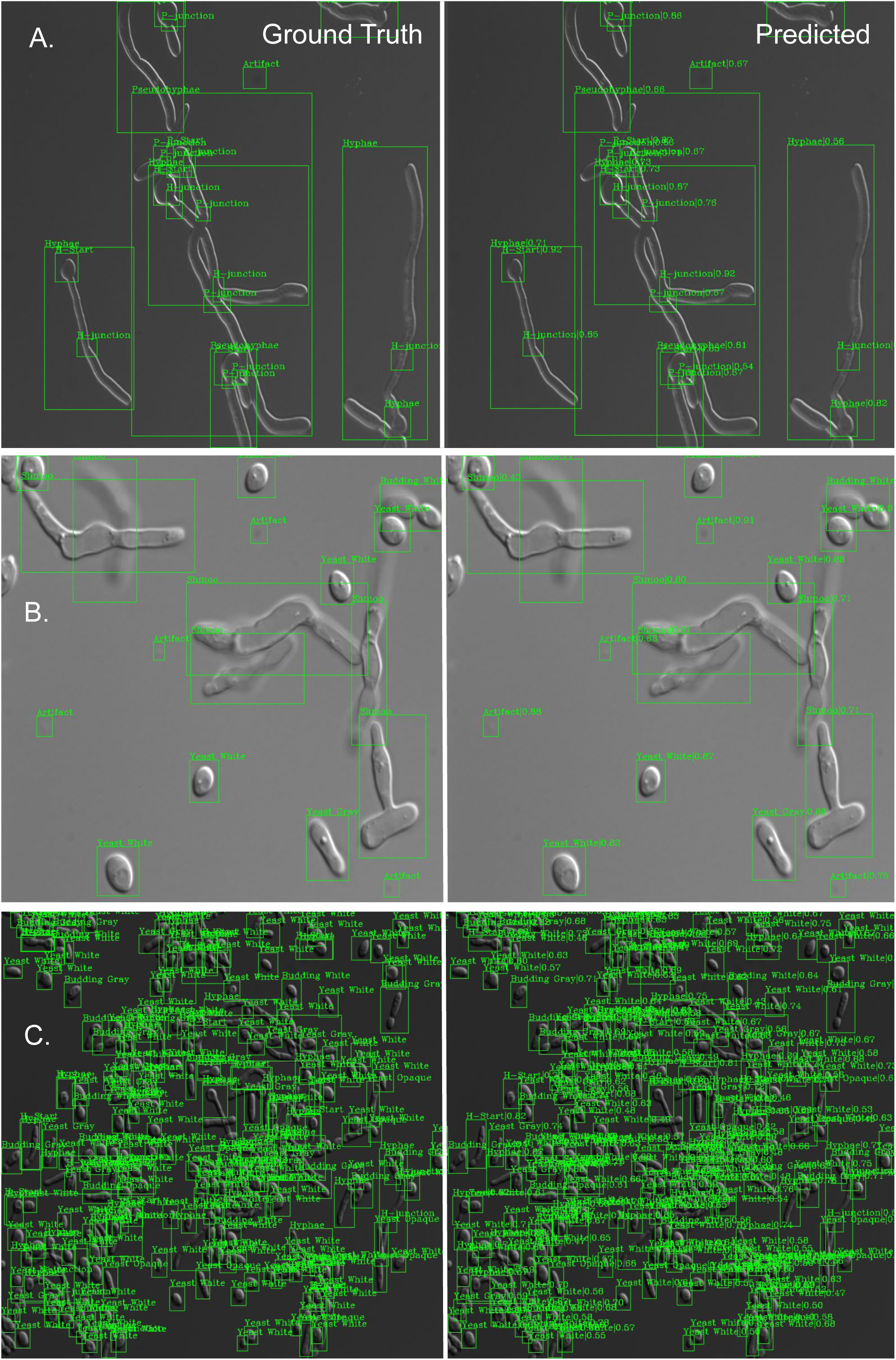
Representative images of the ground truth in the Varasana validation set (left) each of which consists of a bounding box around the object and a class label, and the predictions made by the FCOS Candescence (right) which also have a score between 0 and 1 from the softmax layer of the FCOS representing the strength of belief in the classification. Panel **A** depicts an image we found difficult to label as the objects have both pseudohyphal and hyphal properties. Nevertheless, Candescence recapitulates our classifications. Note that if an object is identified as hyphae, the junctions and start are also labelled as H; this is also true for pseudohyphae. This suggests that Candescence is learning to classify not only on the image but also as a function of the predicted labels of the same object. The image in Panel **B** contains a diverse collection of classes. Note that we labelled gray-like only by their size and “texture” (smaller but rectangular like opaque, and more gaunt than white cells). Overall, these categories witnessed the highest number of classification errors. Candescence predicts well even for highly dense and diverse images such as Panel **C**.

### Candescence almost never hallucinates but has some blind spots

The performance of our classifier can be decomposed into its object detection and object classification components. For object detection, false positives refer to cases where Candescence predicts a bounding box in a location of the image that does not have a ground truth bounding box. Using the 94 images of the validation dataset with an IoU of 0.5 and a range of thresholds for τ, we manually examined all false positive predictions. **Supplemental Figure 3** depicts all 358 such hallucinations for τ = 0.25. It is difficult in some dark images (n=13) to detect an object and these instances were confirmed as true hallucinations by Candescence. In approximately 30 of the remaining cases, there is in fact an object in the bounding buy but the predicted classification is incorrect. Essentially the labellers (missed bounding box) and Candescence (incorrect class) made a mistake. In all remaining 315 cases, Candescence was correct. After adjusting the performance measures by removing the effects of these human errors, the recall, precision and F1 rose to 85.1%, 80.7% and 83.2% respectively (**Table 2A**, bold red, **Methods 5**). **Figure 3A** provides a small sample of correct and incorrect hallucinations.

**Figure 3.**
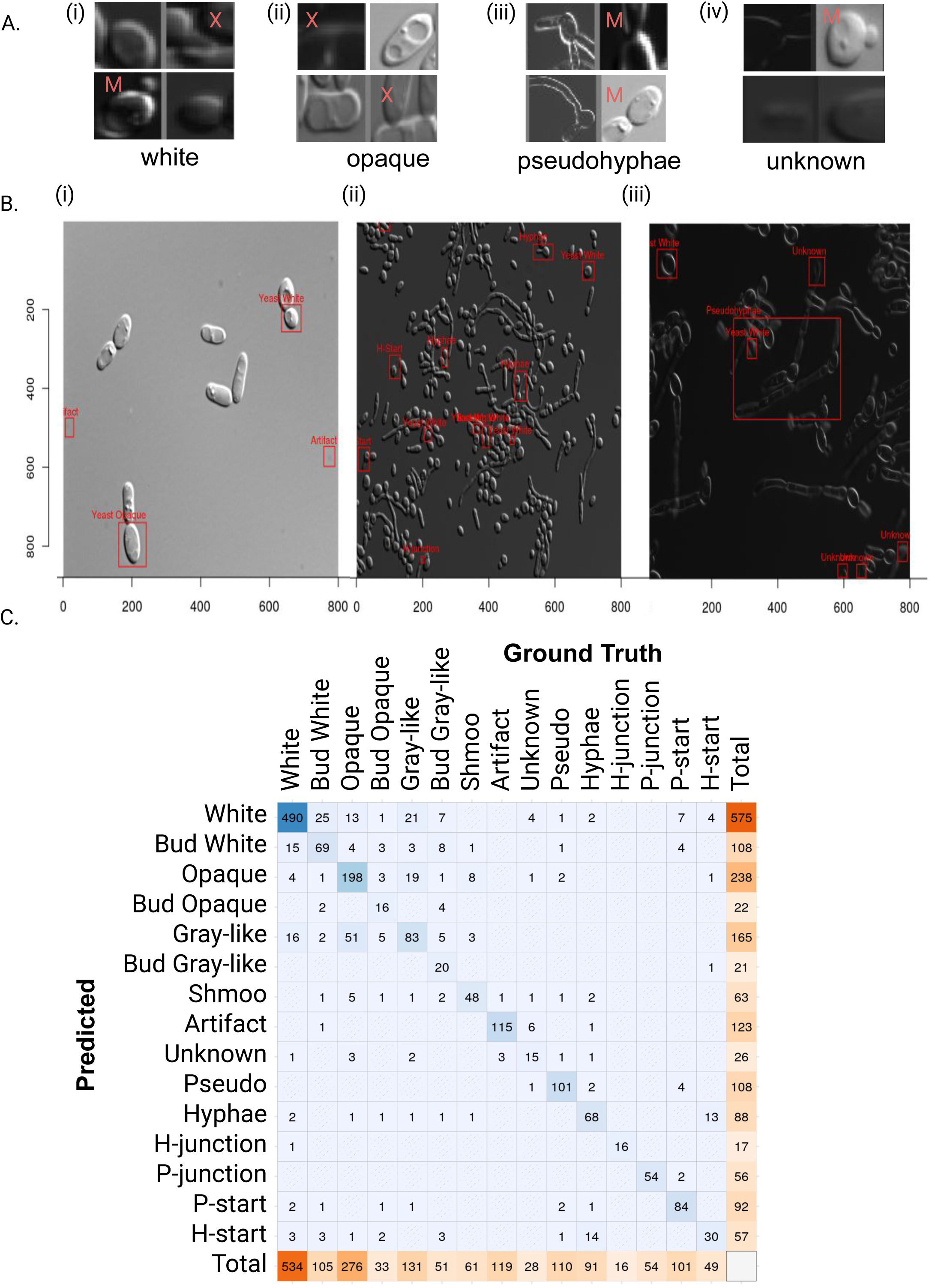
**A.** Representative images chosen from the complete collection in Supplemental Figure 3 of all false positive object identifications from Candescence. This set of hallucinations is subdivided by the class label returned by Candescence. X indicates that we truly are not able to see an object (a true hallucination) and M indicates that Candescence was indeed correct to predict a bounding box at that location (missed by the human annotators) but its subsequent classification disagrees with our criteria. Subimages lacking an annotation indicate that Candescence correctly identified and classified the object, which represents a human error. Panel **B** depicts three partial images with false negatives (“blindspots”) labelled with red boxes. Unannotated cells in **(i-iii)** were correctly handled by Candescence and their bounding boxes have been removed for clarity. **(i)** This is an example of a recurrent problem where the probability from the softmax is distributed over two or more labels (e.g. white and budding white) presumably because it has difficulty to guess whether they are only touching versus still attached. The shared probability causes both to fall below our threshold τ. **(ii)** Although Candescence performs well, blindspots arise presumably due to the dense packing of cells. **(iii)** Our strategy during labelling was to leave cells that were partially outside of the field of view unlabelled. Such edge effects cause some problems and are often predicted as Unknown. Lastly, bounding boxes for large filamentous *C. albicans* are sometimes missed, especially if other cells are within the vicinity. Panel **C** depicts the confusion matrix for Candescence with columns corresponding to ground truth classifications and rows corresponding to Candescence predictions.

Candescence however does miss several bounding boxes in the ground truth dataset (n=301 false negatives). Approximately one-quarter of such blindspots correspond to artifacts. Artifact is a heterogeneous class that was used to label all defects in the microscopy images regardless of their individual visual qualities. As such, it is perhaps understandable that Candescence does not learn to predict this class well as there is no consistent set of attributes. Approximately one-quarter of the blindspots are related to objects labelled as yeast white in the ground truth dataset (**Table 2B**). We observed a recurring pattern across these cases: often the white cell was physically adjacent to a second cell (**Figure 3Bi**). Forensic investigation of the neural network suggests that the softmax function of the FCOS distributes the probability between the yeast white and budding yeast white uniformly. This causes the score for both categories to fall below our chosen threshold τ = 0.25; hence Candescence fails to identify a bounding box in that location of the image. This also occurred between opaque and budding opaque. In fact, more than one third of the images of the validation dataset had at least one problematic prediction of this form.

Candescence failed to identify several white cells in dense images (**Figure 3Bii**), and failed to label several germ tubes cells as H-start. As well, P-junctions in pseudohyphae enriched images were often hard to identify (∼10% of all blindspots). A P-junction looks like two adjacent cells with a bright region in the fluorescent image. This pattern is not unlike countless other locations in the images that capture two adjacent cells, or budding cells. Our hope was that Candescence would learn to associate the presence of P-junctions with the larger bounding box of the pseudohyphae itself along with the “nearby” P-start. There is some indication that Candescence has learnt this calculus, although there are many P-junctions per instance of pseudohyphae and they can be quite distal to the P-start. Lastly, some cells near the edges of the image were overlooked such as the example in **Figure 3B iii**. Overall, many of the blindspots occur in dense, crowded images similar to **Figure 3Bii-iii**.

### Candescence exhibits high classification accuracy

Here we restrict attention to only those objects which have been correctly located in the images (n=2104) and investigate the classification performance of Candescence. In total there are 382 errors (accuracy of 81.8%; **Table 2**). **Figure 3C** depicts the confusion matrix across the 15 classes in the validation dataset with diagonal entries corresponding to correct classifications. Comparing our labels to predictions, we observe pronounced confusion between opaque and gray-like with some 51 opaque cells classified as gray-like, and 19 gray-like as opaque. Significant but slightly reduced confusion exists between white and gray-like. In fact, misclassifications between white, gray-like and opaque account for almost one-third of all classification errors. Although there is considerable confusion between these three classes, we note that the statistical performance is still far better than random, suggesting that our visual inspection and labelling was at least partially consistent. A smaller degree of confusion exists between hyphae and H-start. The softmax appears to diffuse probabilities across H-start, white and budding white, for some bounding boxes. This is perhaps another manifestation of the difficulties we observed with blindspots depicted in Figure 3B.

### Candescence retains its capacity to classify genetically perturbed *C. albicans* in the test set

Our test set of 904 images serves as an independent measure of performance. Some images of SC5314 or SN148a strains without genetic modifications had been left out of both the training and validation set but were grown under the same conditions needed to induce different morphologies. We compared Candescence predictions to manual curation for images referenced in **Table 1A** but differences in performance between the validation and test set were insignificant (comparison of proportions *χ*^2^ test, **Methods 5**).

The test also contains *C. albicans* colonies grown with a variety of conditions, preparation protocols, and genetic perturbations (**Table 1B, Supplemental Table 1**). The four perturbed genes all have well-established and central roles in filamentation processes and the regulation of morphology. Modifications of these genes generated cells that can differ visually from wildtype morphologies, creating an interesting challenge to the ability of Candescence to identify and classify cells.

Our panel includes a transformed SC5314 strain with a CRISPR/Cas9-based homozygous deletion of Unscheduled Meiotic gene Expression (UME6), a Zn(II)2Cys6 (zinc cluster) transcription factor that controls transition to true hyphae by maintaining expression of filament-specific genes in response to inducing conditions. Although cells lacking UME6 are able to form germ tubes, hyphal extension is limited^60^. Our panel also includes Biofilm ReGulatory 1 (BRG1) encoding a transcription factor that recruits the histone deacetylase Hda1 to hyphal-specific promoters and removes Nrg1 inhibition to promote filamentation. Filamentation is decreased in brg1Δ/brg1Δ cells^61^. We did not detect a statistical significant difference in performance of Candescence for both *UME6* null and *BRG 1* null cells under conditions inducing both the white morphology (**Supplemental Figure 4A**) and the pseudohyphae morphology (**Supplemental Figure 4B**).

Regulator of Hyphal Activity 1 (*RHA1*) encodes a zinc cluster transcription factor that serves as a regulator of the Nrg1/Brg1 switch. Hyperactivation of Rha1 can trigger filamentous growth in the absence of external signals or in the presence of serum can bypass the need for Brg1^62^. Loss of Rha1 function leads to reduced ability to generate hyphal growth in the presence of external signatures. Loss of both Rha1 and Ume6 ablates filamentation completely. Although Rha1 null cells generate yeast white cells with standard appearance when grown at 30℃ in YPD, they produce smallish pseudohyphae with reduced branching when grown at 30℃ with serum added to the YPD. Candescence does successfully label pseudohyphal substructures including P-start, P-junction, but there is some increased confusion with hyphae as exemplified in **Supplemental Figure 4C**. A minor depreciation in accuracy was observed and this was statistically significant (p < 0.05, type 20 **Table 1B**).

We cultured *RHA1* gain of function (GOF) mutants constructed using the zinc cluster hyper-activation technique from Schillig and Morschhaeuser^63^ and observed that cells grown at 30°C in YPD formed pseudohyphae that tended to look like wildtype (**Supplemental Figure 4D**). Some of these images were used in the training and validation datasets and performance remains the same on the omitted test images. However, the performance of Candescence is statistically worse in RHA1 GOF/UME6 null cells. Although these cells present a small pseudohyphae morphology, the images tend to contain multiple tightly packed clusters, which we hypothesize contributes to an increase in the number of blindspots (**Supplemental Figure 4E**).

RHA1 GOF/BRG1 null cells have a morphology distinct from wildtype hyphae and pseudohyphae, comprising elongated chains without branching and with less pinching at junctions (**Supplemental Figure 4F**). Here Candescence most often labels these cells as hyphae. Interestingly, if an object has been labelled as hyphae, the junctions are labeled as H-junctions and not P-junctions, although often the characteristic pinching of pseudohyphae is clearly present. This may suggest that the classification rules discovered by the deep learner go beyond appearance and use information regarding the labels of nearby objects.

Candescence again experiences a loss in performance with RHA1 GOF/*BCR1* null cells grown at 30°C in YPD only. Bcr1 is a C2H2 zinc finger transcription factor which regulates a/*α* biofilm formation and cell-surface-associated genes. The homozygous null variant exhibited decreased adhesion, biofilm formation, and cell size. There is conflicting data regarding whether it promotes or inhibits filamentous growth, however cells which do transit to a filamentous morphology appear abnormal^64,65^. In the Varasana image set, this mutant strain generates cells that appear to branch similarly to pseudohyphae but the individual cells are yeast white or budding yeast in appearance. They tend to form thick clusters in the image (**Supplemental Figure 4G**). The performance decrease is highly significant (p < 0.001) with many cells in dense clumps labelled as white or budding white. Candescence does however identify the location of the vast majority of cells and often correctly labels isolated objects as pseudohyphae. When RHA1 GOF/*BCR1* null cells are grown at 37°C with serum added to the medium (types 13, 17, 20, **Table 1B**), we observed large pseudohyphae and Candescence classifies correctly at the same rate as the validation set (**Supplemental Figure 4H**).

### The space of *C. albicans* morphologies is complex and continuous

We observed considerable cell-to-cell morphological variability across the images. Heterogeneity arises due to technical variations (e.g. light intensity, focus), natural biological programs (e.g. cell cycle affecting size/shape), and transitions between morphologies (e.g. growth of a hyphal cell from germ tube). These sources of heterogeneity complicate both the manual labeling procedure during construction of the learning set and the downstream ability of an FCOS to correctly assign morphology. Our goal here is to quantitatively explore the complexity of the Varasana compendium in an unbiased manner.

Towards this end, we developed an unsupervised approach based on a variational autoencoder (VAE)^66^. A VAE is a generative artificial neural network that uses a game theoretic approach involving two players: the encoder and the decoder. Instances of each object (individual cells from our ground truth annotations) are provided to the encoder. The encoder re-represents these images in the latent (hidden) layers of its network. Typically the latent space is much smaller than the size of the original input, forcing the encoder to build succinct models capturing the most salient features of each image. The goal of the decoder is to reconstruct the original image from only the encoder’s latent representation. The encoder and decoder together are penalized according to how much the reconstructed image differs from the original. This cycle is repeated for many epochs across the training and validation sets. Note that this encoding is built in an unsupervised manner as it does not make use of the class labels (morphological assignments). Our particular VAE uses several convolutional layers to encode each image in a 2D latent space across all wildtype (SC5314, SN148a) cells detected by the FCOC-classifier and *β* parameter that down-weights the Kullbach-Leibler component of the loss function (**Methods 6, Supplemental Figure 5**).

After training, the resultant encoder was applied to all cells from the test and validation sets and the two-dimensional latent space visualised (**Figure 4**). Rather than islands of distinct isolated cells, we observe an unbroken continuum in both the V1 and V2 axes. Although some dimensions of the latent space capture specific morphologies (e.g. shmoo, hyphae and pseudohyphae on the left end of the first V1 dimension), there is a ubiquitous imperfect separation between all of the morphologies. This suggests that almost all morphologies have instances that are difficult to differentiate from one another. Yeast white cells span almost the entire first latent dimension, overlapping in some regions heavily with gray-like and opaque. Several technical artifacts including light intensity drive the scatterplot especially in the first V1 dimension. The second V2 dimension primarily captures differences in the size and texture of the cells.

**Figure 4.**
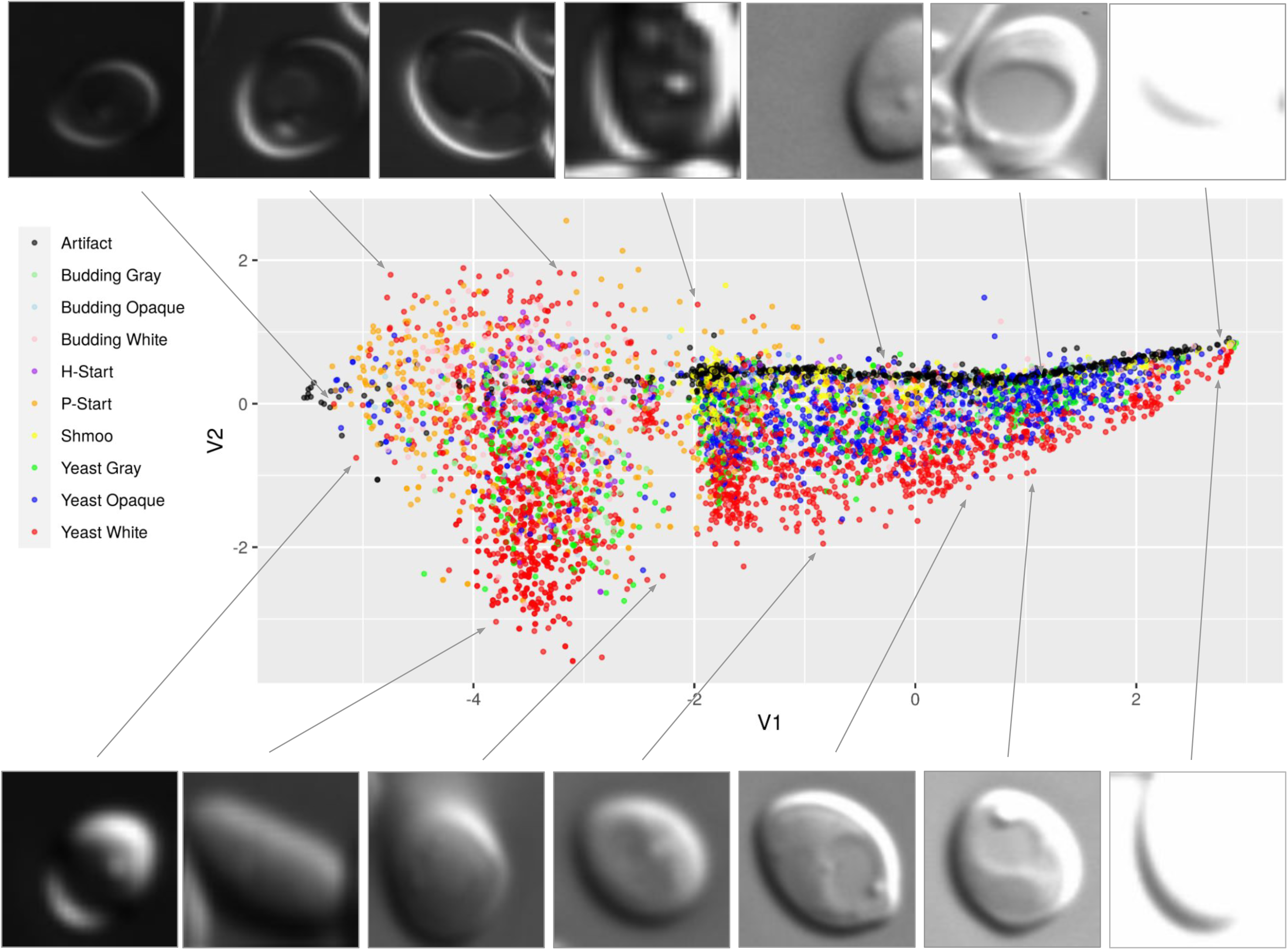
depicts a scatterplot of the two dimensional latent space of the VAE on the training and validation datasets. The junction, hyphae, pseudohyphae, and unknown class have been removed from the training and visualization of the VAE. The V1 dimension of the VAE is strongly associated with light intensity of the image. The vertical V2 dimension captures other variability primarily related to size and texture of the cells.

### Capturing the canonical forms of *C. albicans* morphologies: generative adversarial networks

A sufficiently large collection of images will likely capture snapshots of cells in all technical (e.g. different light intensities), developmental (e.g. across each step of the cell cycle) and morphological states (e.g. along the transition from germ tube to hyphae). Our goal here is to build pseudotime models from these images that capture the progression of these technical and biological variables. Our approach is based on generative adversarial networks (GANs). Intuitively, this deep learning technique re-represents the images in a latent space in a manner that captures this step-by-step progression. Then computational techniques can search for trajectories through the latent space that correspond to a specific effect of interest (e.g. a specific transformation between two morphologies). This allows us to derive continuous “movie-like” models of cells morphing along this trajectory.

GANs are trained using a game-theoretic adversarial approach that pits two deep networks - the *generator* and the *discriminator* - against each other^67^. The purpose of the generator model is to create “fake” examples of *C. albicans* cells with different morphologies. These fake images are created to deceive its adversary, the discriminator model. The generator is allowed to feed both fake and real images to the discriminator model, whose goal is to differentiate between the two. The generator is then told which images the discriminator got right or wrong; this information is used to update the generative model (which corresponds to updating parameters of the neural network). This creates an “arms race” between the deep networks. After a sufficient number of epochs, the generator will ideally produce fake images of *C. albicans* morphologies which cannot be distinguished from true images by the discriminator. Our GAN is based on the computationally accessible method from Liu and colleagues^68^ and learnt using the training and validation components of the Varasana dataset (**Methods 7**).

After training, we exploit the generator to explore trajectories in our latent space. We start with two images representing the end-points of a process of interest. For example, the left end-point might be a real image of a yeast white cell while the right end-point might correspond to an opaque cell. We then used a process called inversion^69^ to find a representation of these two images in the latent space of the generator (**Methods 7**). **Figure 5A** depicts examples of target images and their nearest neighbour in the latent space. Next, the system finds a linear path between these two points in the latent space, so that the nearest neighbour of the final “fake” image at *t*_7_ is the real image at the right endpoint. Lastly, the intermediate “fake” images are reconstructed from the latent space to provide a visualization of the trajectory using the generator function. **Figure 5B (i)** depicts the results of applying this procedure to find yeast white to opaque morphological switch. Interestingly, the system seems to arrive at a decision point at *t*_7_. At this point, the trajectory could bifurcate towards budding white, although in this case it continues towards opaque. As a comparison, when we asked for a trajectory from yeast white to budding white (**Figure 5B ii**), the trajectory stays true to the yeast form and does not appear to wander towards the opaque morphology. The hypothesized bud is perhaps somewhat disproportionately large compared to its mother, and larger than the bud of the real image. Panel **5B (iii)** presents a trajectory that starts with a budding opaque cell. Both cells appear to bud a second time from *t*_4_ through *t*_6_. The progeny of the daughter cell (left) disappears from the trajectory. Although imperfect, the system appears to have learnt a reasonable model of pseudohyphal development from relatively few (∼200) images. Continuous movies for each of these trajectories are available in the supplementary material.

**Figure 5.**
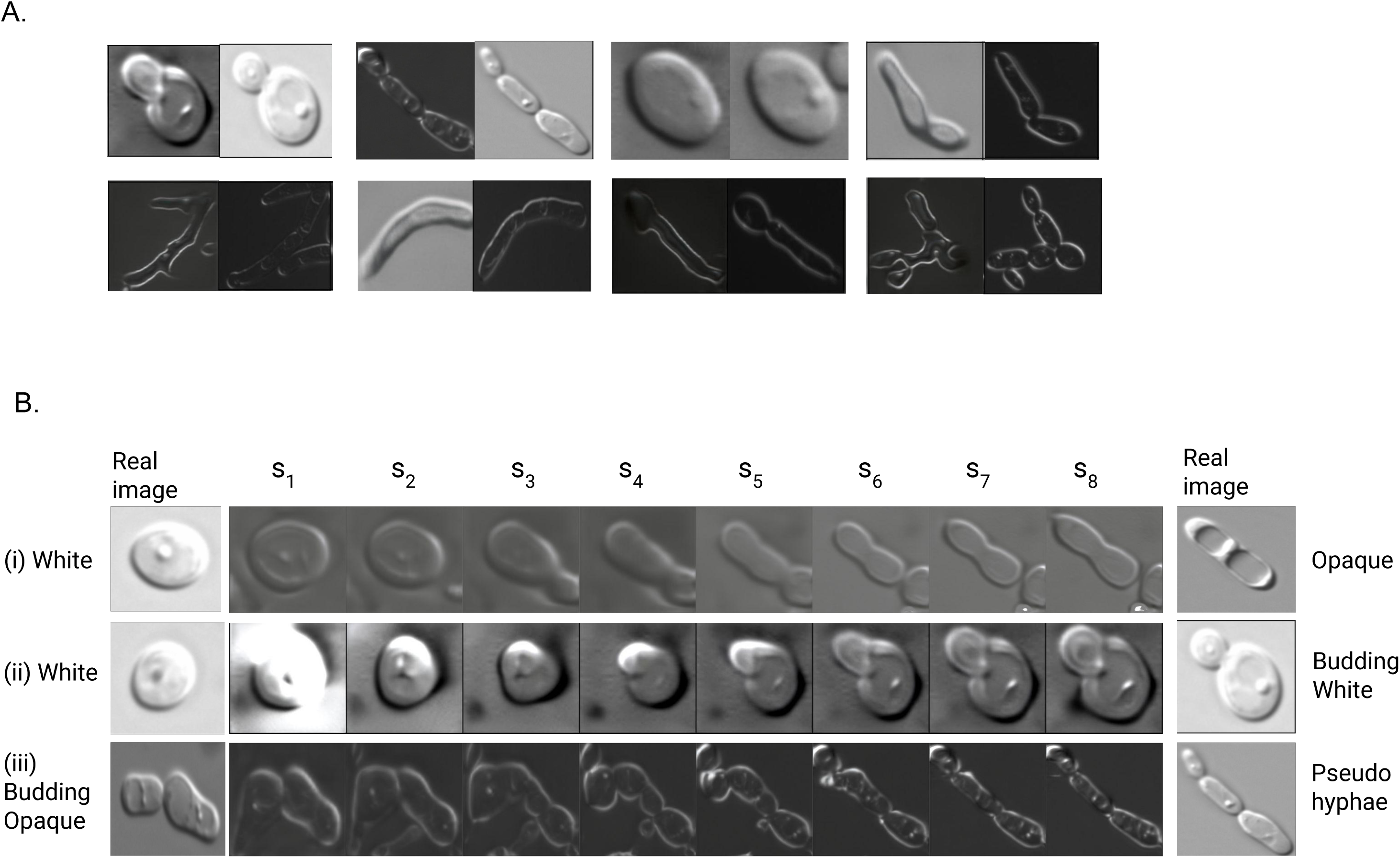
**A.** For each pair of images, the left is a synthesized image produced by our model, and the right image is the image found from the real training data that is its nearest neighbour under the LPIPS metric. **B.** Three separate trajectories. Here the left and right most images correspond to real images from the training data. The sequence of images at points *s*_1_ to *s*_8_ are produced by a linear interpolation between the real images in the generator’s latent space in the GAN.

### Detecting deviations from standard *C. albicans* morphologies: anomaly detection with GANs

Microscopy is routinely used to judge whether a specific genetic or environmental perturbation has led to an observable phenotype. This could manifest as a visual change in the composition of cells in an image (e.g. an increase in opaque cells versus control), a difference in the spatial distribution of cells in the image (e.g. clumping of cells), or a change in appearance of the cell (e.g. shortened hyphae, large vacuoles). In this section, we build upon our GAN-based morphology models to address the third issue, namely a system capable of deciding whether the cells in an image deviate significantly from the space of wildtype *C. albicans* morphologies. Our hope is that the deep learner is more sensitive than “eyeballing” microscopy images when manually attempting to investigate if an experimental strain is abnormal.

The algorithm starts by mapping a target bounding box (representing a single cell, hyphae or pseudohyphae) into the latent space of the GAN’s generator (**Methods 8**). The nearest neighbour in the latent space is found; this corresponds to the object from the training and validation dataset that is visually most similar to the target. The distance between the target cell and nearest neighbour is computed using a specialized similarity metric developed from the Learned Perceptual Image Patch Similarity^70^ measure. The intuition is that distances between a target with a normal morphology will have a nearest neighbour that is closer in the latent space than a target with a very abnormal morphology.

**Figure 6** represents an example of using this test with a *RHA1* GOF/*BCR1* null colony. Candescence is first used to identify the objects and their morphology in the image, and the anomaly score is computed for each such object in the latent space built from the Varsana learning set. When we compare the distribution of anomaly scores between all images from this mutant colony (type 13) and compare them against a collection of “wildtype” pseudohyphal cells (type 65), we observe a statistical enrichment of outliers with abnormal morphology (Kolmogorov-Smirnoff test, p < 0.01). We generally do not observe differences at the low end of anomaly scores, as almost all images from genetically perturbed colonies still contain many examples of cells with normal morphology.

**Figure 6.**
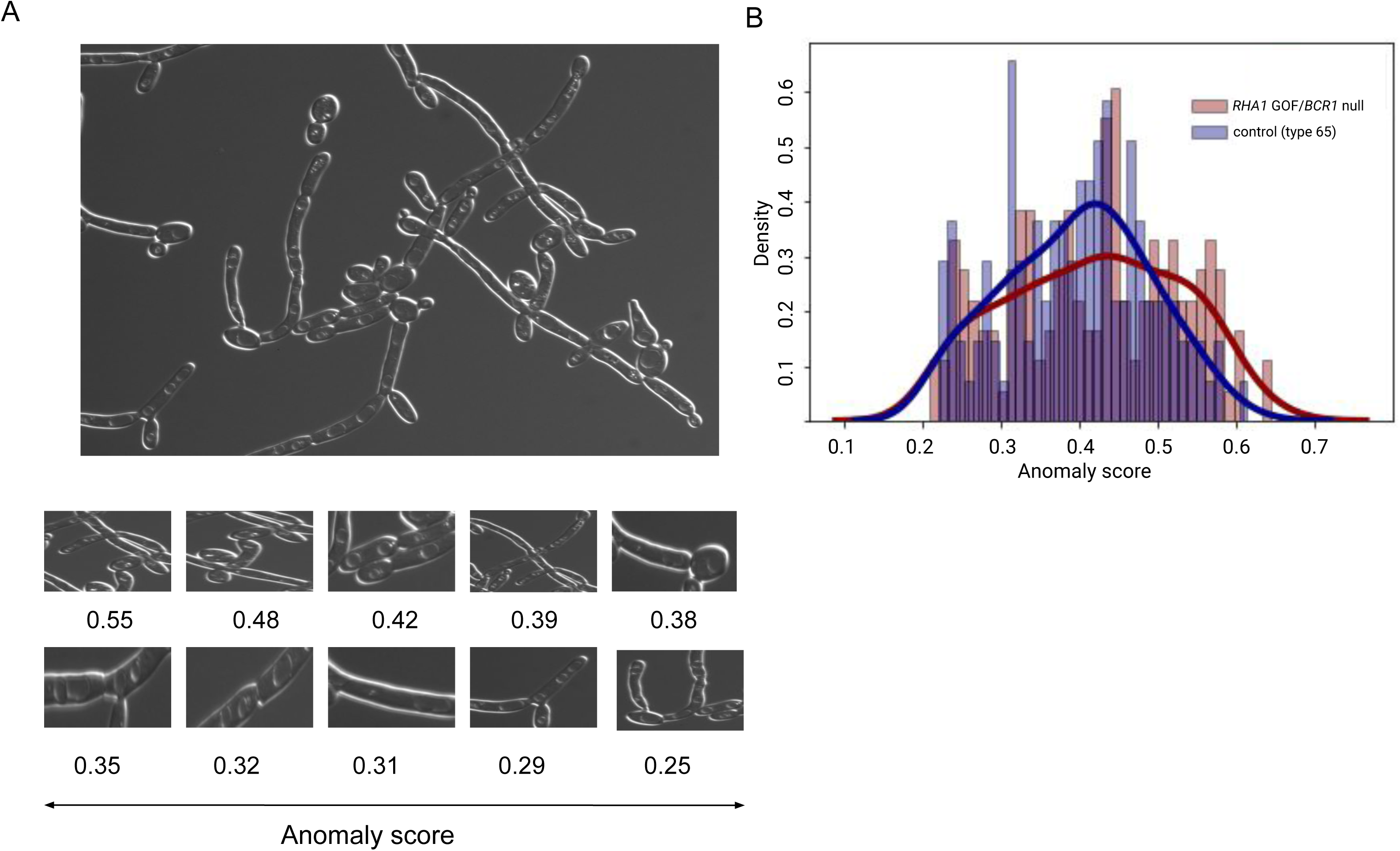
**A.** Example of anomaly detection using a *RHA1* GOF/*BCR1* null colony (UID 230 of type 13). Panel **B** enlarges a series of bounding boxes predicted by Candescence with their associated anomaly score. The histogram of panel **C** compares the distribution of anomaly scores between all images of type 13 versus type 30, a collection of normal appearing pseudohyphal and hyphal cells. There is a small but statistically significant enrichment of cells from the *RHA1* GOF/*BCR1* null colony with elevated anomaly score differences (Kolmogorov-Smirnoff test, p < 0.01).

## Conclusions

Candescence includes a multi-object detection algorithm capable of accurately classifying nine *C. albicans* morphologies. It is based on a fully convolutional one-stage (FCOS) architecture which both locates objects and classifies them with high accuracy. The system is trained with the Varasana learning set consisting of ∼1,200 total images. The training and validation datasets consist of 310 images which have been manually annotated with bounding boxes and class assignments. Using transfer learning, the starting point for training is ResNet-101, a network capable of locating and classifying common household items and pets. It is possible that the image building blocks (textures, edges, colours) encoded in ResNet-101 are not optimal for (DIC) microscopy. As our collection of *C. albicans* images grows, it may be feasible to build a microscopy-specific analog of ResNet-101 and improve performance.

Since a flat-structured learning set led to suboptimal performance, we developed a six grade curriculum learning set and ordered examples by increasing difficulty. To the best of our knowledge we are the first to use a cumulative strategy where images appear at some grade and re-appear in all subsequent grades. We hypothesize that this approach removes the need for layered freezing strategies. This is advantageous, since optimization of hyper-parameters is computationally expensive and the space of possible freezing strategies grows exponentially in the number of grades and network layers.

Candescence has very good performance. Given that an object has been correctly regressed in the image, it will be assigned the correct class label with ∼82% probability. Misclassifications tend to occur between morphologies that ‘overlap’ as highlighted by the VAE plots of **Figure 4**. Confusion tends to exist between vegetative morphologies and their budding/mating forms (e.g. from white yeast to budding white). The difficulty here for Candescence is to judge cases where the daughter cell is large but still attached versus detached but still physically adjacent to its mother. We hypothesize that a larger set of training examples will remove this confusion. Inclusion of time lapse images of specific biological processes would also improve performance. Moreover, integration of the matched fluorescent images with the current grayscale DIC image would perhaps provide the learning procedure with the necessary information to judge whether separation has occurred. It would be straightforward to encode the fluorescent image as an added dimension beyond the gray scale 800×800 input currently used. We did not pursue this avenue in the first version due to complications in the training procedure, since some fluorescent images were not available.

The construction of any learning set requires consistent labelling rules. In our setting, there were visible differences in sizable subpopulations that stretch from yeast white to opaque. **Figure 4** reinforces that this heterogeneity is continuous with several areas enriched for cells with a common but perhaps non-canonical appearance. This includes a large number of small rectangular “gaunt” cells we labelled as gray-like. Candescence is at times confused between the yeast white, gray-like and opaque morphologies, an expected observation given that there were many images for which we were unable to form consensus as a group. Absolute assessment of true gray versus our gray-like is out of reach for this study as we lack the necessary molecular markers as per the original findings of Tao and colleagues^16^. Although false classifications are enriched between white, opaque and gray-like, the success rate is still very good, suggesting that this dichotomy does exist in the images. If these cells had instead been randomly assigned the three class labels without regards to physical appearance, it is highly unlikely a classifier could learn to distinguish them with statistically significant performance. Although the relationship to the Tian et al. gray cells remains unsettled, it does suggest that there is interesting substructure across the vegetative forms that is perhaps not captured by our current dichotomy; other cryptic morphologies could exist. These physically distinct subcolonies could represent simply canonical vegetative forms at specific moments of their development, or could represent phenotype diversity that arises in response to an environmental or communal cue^71^. Building upon this primeval version, Candescence may eventually allow for the interrogation of community structure and interaction.

Imagery of stained or fluorescent reporter molecules would help resolve issues of cryptic morphologies and would extend the capacity of the system to classify using subcellular features. This technique has been used successfully to explore changes to *S. cerevisiae* in the presence of genetic perturbations^55^, although Baker’s yeast does not have as large a range of morphologies as *C. albicans*.

Our system initially appeared to have significant challenges during the object detection step. However, careful analysis of the false positive detections suggest that many such events are in fact not “hallucinations” but “false false positives”. That is, they correspond to true events in the image files that we missed during the manual annotation procedure. A large portion of these events correspond to either subtle technical artifacts or small cells in crowded images.

Candescence appears to have some blind spots, missing cells that are annotated in the ground truth dataset. The tradeoff between false negatives and positives is controlled by underlying IoU and threshold **τ** parameters. We observed that accurate bounding boxes were indeed regressed for the majority of these false negative objects but such objects required an abatement of either the IoU or **τ** parameters before they were reported as positive predictions. Our forensic analysis of the neural network suggests that this is due to confusion between, for example, yeast white and yeast budding white; the probability is amortized over two or more classes and therefore drops below **τ**.

Furthermore with respect to blindspots, we hypothesize that the differences in size and shape between the morphologies (e.g. yeast white versus pseudohyphae) induce different distributions of IoU and **τ** scores. Therefore, in images containing diverse cell types, the single universal **τ** is essentially too conservative for some morphologies but too liberal for others. It is an interesting future challenge to modify FCOS-based classifiers to adjust for this heterogeneity in a statistically sound manner.

We explored in the independent test set a range of genetically altered *C. albicans* populations involving genes *RHA1*, *UME6*, *BCR1* and *BGR1* with established roles controlling filamentation. The observed changes in classification accuracy is consistent with the fact that these morphologies represent shifts away from wildtype forms. The Candescence response to these perturbations is intuitive and largely retains its performance.

Variational autoencoders (VAEs) provide a convenient tool for modelling the diversity of cells caused by natural cellular programs, morphologies and technical artifacts. We observe a nearly unbroken, continuous distribution of points in our two-dimensional embedding, suggesting that the underlying space of *C. albicans* morphologies are also continuous and overlapping, as perhaps expected. We certainly cannot rule out the possibility that a more advanced architecture for the VAE would better separate the morphologies. However we stress that this specific VAE easily separates the vast majority of points in other deep learning sets including MNIST^72^ and Omniglot^73^, suggesting the *C. albicans* morphology is at least as difficult as these well-studied learning challenges.

To the best of our knowledge, this is the first attempt to capture the space of C. albicans morphology in a continuous manner that respects technical variation, developmental processes and morphological transitions. Our intention is that the GAN models can be used to automatically detect new morphologies that are perhaps subtly different from our current dichotomy. Using tools for detecting anomalies via the generator’s latent space, we show how cells displaying non-canonical morphological forms can be flagged and quantified. This should find practical value in microscopy based studies: the tool will provide the community with a central resource that not only houses all *C. albicans* images but also unbiased models developed from those images that extended to non-canonical morphologies. Future studies will benefit from the ability to compare their images across this synthesized sum of knowledge. The technique should be straightforward to transfer to other fungi.

We have shown that the deep learning-based approaches are able to recapitulate the classification rules that are encoded by our choice of labelling strategy. It is unlikely that our labelling strategy is correct and other labellers with more or different expertise may have chosen a different way to partition the learning set and assign labels to individual examples. Although we attempted to be as consistent as possible when assigning class labels in Varasana version 1.0, our labels are certainly imperfect and open to debate. The classifier does however function far beyond random guesses, suggesting that our scheme has some value.

There is evidence that Candescence is able to “overcome” errors and inconsistencies between labellers. This is a well documented problem in image recognition research including computational pathology^74^. We argue that the computational techniques introduced to computational pathology are largely applicable to fungal systems. For example, machine learning based analysis of medical images has been shown to be more sensitive than trained pathologists identifying small events and complex patterns beyond perhaps human capacities^75^. As Candescence evolves to include stainings, fluorescent markers and a greater spectrum of microscopy imaging techniques, it may reveal cryptic cell or subcellular structure or community organization. Computational approaches in imaging provide a means to combine different modes of data, and to provide downstream analyses that integrate this information in a statistically sound manner^76–78^. For *C. albicans*, this might entail the integration of information concerning strain, growth conditions and genetic challenges together with images to better understand the composition and dynamics of colonies. Imaging standards analogous to the Digital Imaging and Communication (DICOM)^79^ for microbial systems including imaging of host tissue would enable better data sharing across the fungal community and allow for “hive analysis”^80^.

Last and perhaps most importantly, a surprising degree of disagreement has been observed (and quantified via Cohen’s ***κ*** score) between expert pathologists when challenged with the same images^81^. Our limited experience with *C. albicans* morphology suggests that there may be similar disagreements across experts in this field. Such differences may hint at important alternative classification schemes. These differences may be important as Candescence is extended into clinical samples where the presence of *C. albicans* and its morphology are considered concomitantly within their host tissue. Our effort here represents an opportunity for the community to kernelize their knowledge of the dynamics of morphologies in a quantitative objective manner.

## Materials and Methods

### 1. A fully convolutional one-stage object detector for morphology classification

There are fundamentally two computational problems underlying multi-class, multi-object detection. The first problem is to detect the locations in an image where objects exist; this is a regression to determine the four coordinates corresponding to the corners of the bounding box. In our case, the number of objects per image ranges up to ∼100 (**Supplemental Figure 1**) and objects may overlap in these images especially with respect to the filamentous morphology. Most object detectors rely on pre-defined anchor boxes. An anchor box is a rectangle that bounds an object in an image. Approaches that use anchor boxes make educated guesses where objects might exist in the image in addition to guesses regarding the size, aspect ratio and number of such boxes. The fact that there are exponentially many potential anchor boxes in any image makes this a computationally demanding exercise. Moreover, there are many parameters (size, aspect ratio and number of boxes) that require optimization and re-design on new datasets. The second problem is then to correctly label each object by its class (morphology, start or junction attribute, unknown or artifact).

Here we have opted to use a fully convolutional one-stage object detector (FCOS) for classifying *C.albicans* morphology^56^. FCOSs represent an anchor box-free reformulation of object detection. This is achieved by predicting, for each point in each feature map, the offset position to the top-left and bottom-right coordinates of a bounding box. Five feature maps are produced and each such map is limited to predicting bounding boxes of a predetermined size. For example, the first feature map predicts bounding boxes with maximum area of 30×30 pixels, while the final feature map is used to predict bounding boxes with maximum area of 128×128 pixels. A standard convolutional neural network is used with a softmax head for the classification component. We developed Candescence on top of the open-source implementation of FCOS provided by the machine vision platform MMDetection^82^ (**Figure 1F**).

### 2. Strains and Media

Images of yeast white, opaque and shmoo morphologies were acquired from *C. albicans* SN148a cells grown on YPD agar (1% yeast extract, 2% bacto-peptone, 2% D-glucose, 2% agar and 50 µg/mL uridine). Opaque switching was induced by growing cells on SC Glucosamine media (0.67% yeast nitrogen base lacking amino acids, 0.15% complete amino acid mixture, 2% agar, 1.25% N-acetylglucosamine (GlcNAc), 100 µg/mL uridine). We used 5 µg/mL phloxine B to stain opaque colonies. The shmoo morphology was induced by treating opaque cells with 10 µg/mL α-pheromone for 24 hrs in room temperature shaking at 220rpm.

To Induce filamentation in wild type SC5314, two colonies of cells was grown separately in 5ml of glucose-phosphate-proline (GPP) media^83^, (2.5 mM KH2PO4 (pH 6.5), 10.2 mM L-proline, 2.6 mM N-acetyl-D-glucosamine and 3 mM MgSO47H2O, 20% glucose) for 12 to 16 hours in 30 and 37℃ shaker incubator. The next day, 1 ml of cells from each colony were washed twice with 1ml 1X PBS. Different genetic variants of the SC5314 strain were also used to generate colonies enriched for filamentous morphologies. After growth, cells were washed with 1X PBS and diluted to different OD600 values in fresh liquid Spider, YPD or combinations of YPD and Fetal Bovine serum with or without centrifugation as per **Supplemental Table 1**, which provides a complete list of strains, conditions and protocols.

### 3. Microscopy

*C. albicans* colonies were mounted on slides and stained with a concentration of 2 μg/ml calcofluor white for 20 minutes before imaging. Images of *C. albicans* were captured using a Leica DM6000 upright microscope equipped with 100x (NA 1.3), 60x (NA 1.4) and 40x (NA 0.75) lenses and a Hamamatsu Orca ER camera. For DIC images, samples were captured using DIC optics and the built-in transmitted illuminator of the microscope. For cells labelled with fluorescent probes, samples were illuminated with a 100W mercury bulb (Osram) and passed through filter cubes optimised for illumination of calcofluor white-labelled samples (ex 377/50, em 447/60). We used the fluorescent images during the manual labelling procedure to help decide the best morphological assignment. This was particularly relevant to distinguish between white/opaque/gray-like and their budding variants, and also between junction types for the filamentous morphologies as bud scars are clearly visible. However the fluorescent images were not submitted to the FCOS (or other deep learning tool) during training. They are available as part of the Varasana learning set.

### 4. Image annotation and development of the learning set

In general, all computations were done using Python version 3.7 and R version 3.6.3. Our learning set was prepared using Labelbox (https://labelbox.com), software that facilitates the distributed annotation of image files. As a group, we labelled images following the guidelines from Sudbery et al.^21^, Whiteway and Bachewich^11^, Noble et al.^23^ and Tao et al.^16^. Labelbox assigns images in a manner that guarantees the same image is scored by multiple labellers.

One labeller (MH) modified assignments after the first round of labelling to ensure consistency as best possible across the labellers. A second round of quality control was performed by MH after the grid search was completed and our best classifier identified. Here all false positives and negatives were examined and a decision was made as to whether the instance was a labelling mistake, or a mistake made by the classifier.

### 5. Measures of system performance

Throughout the following TP, TN, FP, FN denote true positives, true negatives, false positives and false negatives respectively. Recall (a.k.a. sensitivity) measures the rate of false negatives whereas precision measures the rate of false positives:

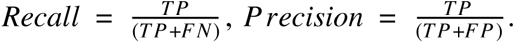

The *F*_1_ measure is convenient as it combines both the recall and precision into a single summary statistic:

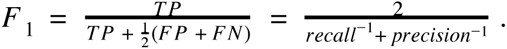

In multi-object/multi-class problems there are two fundamental parameters. The first parameter is related to the object detection component of the FCOS and is termed the Intersection over Union (IoU) value:

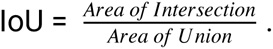

Throughout this manuscript, we use a threshold of 0.5 for the IOU. The IoU controls how closely the predicted bounding box must overlap with the ground truth bounding box to be considered a positive. More stringent IOUs tend to decrease the recall of the system significantly with recall dropping due to a rapid increase in the number of false negatives^84^.

The second parameter is related to the classification component of the FCOS. Here τ represents the minimum value from the softmax of the classification head of the FCOS that must be exceeded if an object is to be assigned a class. More precisely, the score that an object belongs to each of the 15 classes is computed. It is assigned a class if and only if (i) the class has the highest score and (ii) the score exceeds τ.

The dual nature of object detection/object classification problems requires refinement of these fundamental concepts. We say that an object in an image is a TP if and only if the bounding box is predicted correctly (i.e. the IoU > 0.5) and the object is then classified correctly:

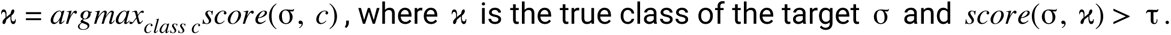

The concept of a true negative in this setting is tricky, since any pixel, which does not belong to a bounding box in an image, is in essence a TN. An object is a FP if and only if either (i) the predicted bounding box does not overlap sufficiently with a ground truth bounding box, or (ii) the predicted bounding box does overlap sufficiently with a ground truth bounding box but the classification is incorrect. Type (i) is termed as *hallucination*. Type (ii) is a misclassification. An object is a FN if and only if we fail to predict a bounding box with sufficient overlap with a ground truth bounding box. We term these FNs *blindspots*.

The mean average precision (mAP) is the standard and preferred approach for measuring the performance of multi-object/multi-class problems^59^. The mAP computes the average of the average precision over τ. Here the average precision corresponds to the area under the precision recall curve induce by a specific threshold for the IoU and varying τ. The maximum value for the mAP is 1.

Within an FCOS, total loss is computed as the sum of three individual losses: center-ness, bounding box and classification loss (**Supplemental Figure 2**). The concept of center-ness loss is specific to the FCOS and represents a mechanism to avoid the identification of multiple, spurious bounding boxes for a single object. It converges to a value of 0.57^56,85^. Bounding box loss measures disagreement between the ground truth location of bounding boxes with the regression produced by the deep learner. Finally, classification loss measures how well correctly identified objects are assigned classes.

We searched the hyperparameter space defined by the cross-product of different settings for the learning rate, momentum, decay, number of epochs per grade, IoU, threshold **τ** and various freezing rates (**Figure 1H**) where boldface font denotes the choice of parameters for the final FCOS. Throughout these experiments, the 800 × 800 input images were subjected to augmentation (random multi-scale flipping). Other parameters included 1000 warmup iterations, a warmup ratio of ⅓ and an SGD optimizer with grad clipping. The mAP and three loss functions described above were used to initially judge the quality of the classifier. The code from running the FCOS and interpreting results is depicted in **Supplemental Figure 7**.

When investigating differences in performance of a classifier between the test and validation sets, we manually curated a subset of test images for each entry in **Table 1A and B** in a manner to ensure that at least 100 cells were labelled (with the exception of the first two entries of Table 1A where there were too few cells). Then we built a contingency table where rows correspond to correct and incorrect predictions, and columns correspond to the test and validation dataset. A χ^2^ test was used with the null hypothesis of no difference between the overall number of correct predictions in the test and validation sets.

### 6. Variational autoencoder (VAE) for unsupervised analyses

The variational autoencoder was constructed using the Keras for R (version 2.4) package^86^. **Supplemental Figure 5** depicts the architecture of the model. Briefly here, the input to the network is a 128 × 128 pixel image. The image is subjected to a series of convolutional layers with a 5 × 5 kernels, 64 filters and strides that successively transformation the representation, followed by a flattening operation that reshapes the representation into a 262, 144 real vector, before a final reduction to a two dimensional latent space. Layers for *z* mean, the *z log* score and for decoding all follow standard VAE procedures (see code for details). Training used the wildtype Varasana training and validation sets in batch sizes of 100 across 20 epochs. The test set objects, which include the genetically modified *C. albicans* variants from Varasana, were not used in training. In particular, we extracted each (ground truth) bounding box across these files and reshaped the images to size 128 × 128. We used a β=0.4 parameter to down-weight the Kullbach-Leibler portion of the loss function^87^. It is straightforward to visualize the resultant two dimensional latent space with a scatterplot.

### 7. Generative Adversarial Networks

We follow the approach from Liu et al.^68^ to build unconditional GAN images. This approach requires computational resources that are accessible to most labs and requires few training samples, a feature important for our setting here. The model is particularly amenable to the disentanglement procedures utilized below. **Supplemental Figure 6A, B** depict the structure of the Generator and Discriminator respectively. We extended the Pytorch-based code available from the authors^88^. The input used for training corresponds to the ground truth bounding boxes of the training and validation datasets, reshaped as 128 × 128 image as was done for the VAE described in Methods 6. Here we used a shift predictor and deformator learning rate of 0.0001, with 2000 steps in batches of size 4.

We follow the framework of Creswell and Bharath^69^ to build trajectories between two (real) target images. Here however we opted to use the Learned Perceptual Image Patch Similarity (LPIPS) metric as the loss function^70^. The core idea is to find both “real” target images in the Generator’s latent space. This requires a so-called inversion which we perform using Algorithm INFER of Creswell and Bharath. Then we use linear interpolation between these two points in the latent space and reconstruct visual representations at user defined points along this path.

### 8. Anomaly detection

Anomaly detection proceeds using the framework from Schlegl et al.^89^ but replacing their residual and discriminator loss with the Learned Perceptual Image Patch Similarity (LPIPS) metric for estimating the similarity between two images^70^. Our approach proceeds as follows.

1. a. Each object in a image file is bounded manually, or; b. Candescence is used to automatically regress bounding boxes for the objects in the target image file.
2. For each target object (that is, for each bounding box or patch) *T*, we find its optimal inversion *z* in the latent space of the generator G using the *INFER*(*T*, *G*) algorithm from Creswell and Bharath modified to use the LPIPS similarity metric. Here *G*(*z*) is a synthetic image.
3. Find the nearest real neighbour *d* of *G*(*z*) across all images *d* in the training set *D* under the LPIPS metric.
4. The anomaly score is then defined as follows:

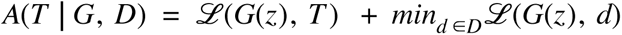

where *D* is the set of training images and ℒ is the LPIPS function between two images.

With respect to 1b, we remark that the performance of Candescence was not severely reduced when given an image with non-canonical morphologies and in this setting the classification returned by the FCOS is not used. Therefore, this automated approach should suffice in most scenarios, unless the change in morphology is very large. However, when the change in morphology is very obvious, we will not need a sensitive algorithm to detect it.

## Author Contributions

VB developed the deep learning algorithm, performed experiments and helped prepare the manuscript. ACBPC, RPO and SM constructed the library of microscopy images used for training. EK and SS developed the Varasana learning set from these images. CL provided expertise in microscopy and analysis of the images. VD, MW and MTH prepared the manuscript. MTH designed the study and obtained funding for the project.

## Data Availability and Reproducibility

The Varasana dataset, code, trained models, supplemental movies and a Jupyter notebook to run Candescence is available from the Open Science Framework at https://osf.io/qdxbp.

---

**Supplemental Figure 1.**
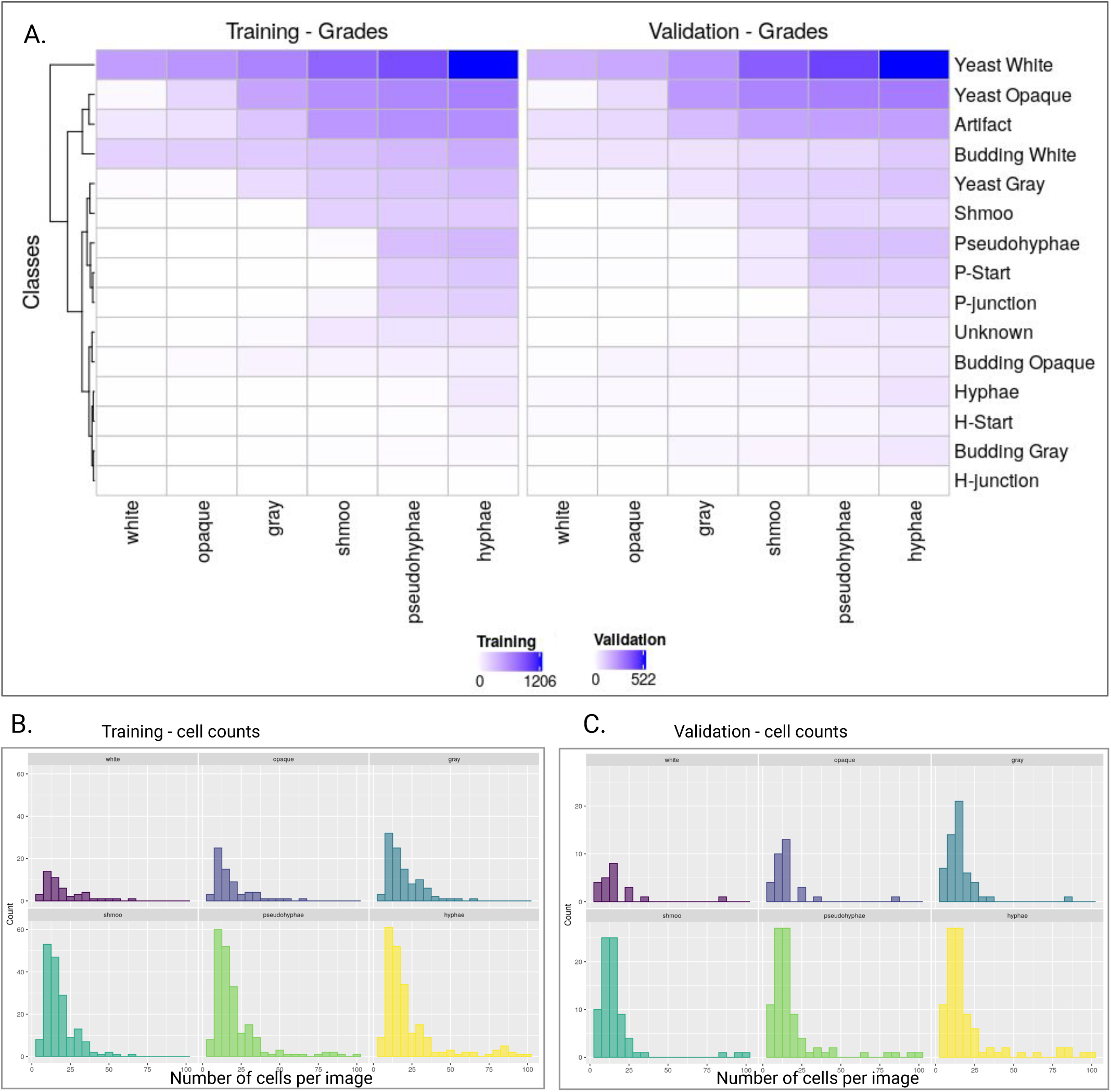
Description of the cumulative curriculum learning set. Panel **A** depicts the frequency of the different classes across the six grades in both training and validation. An image is part of all subsequent grades once it appears for the first time. Images are included in the training and validation sets at a ratio of 7:3. Panel **B** provides histograms of the number of objects (cells, junctions, unknown, artifacts) per image across all six grades. The maximum number of objects in any image was 97, although we note that several test set images exceeded this bound (not shown here).

**Supplemental Figure 2.**
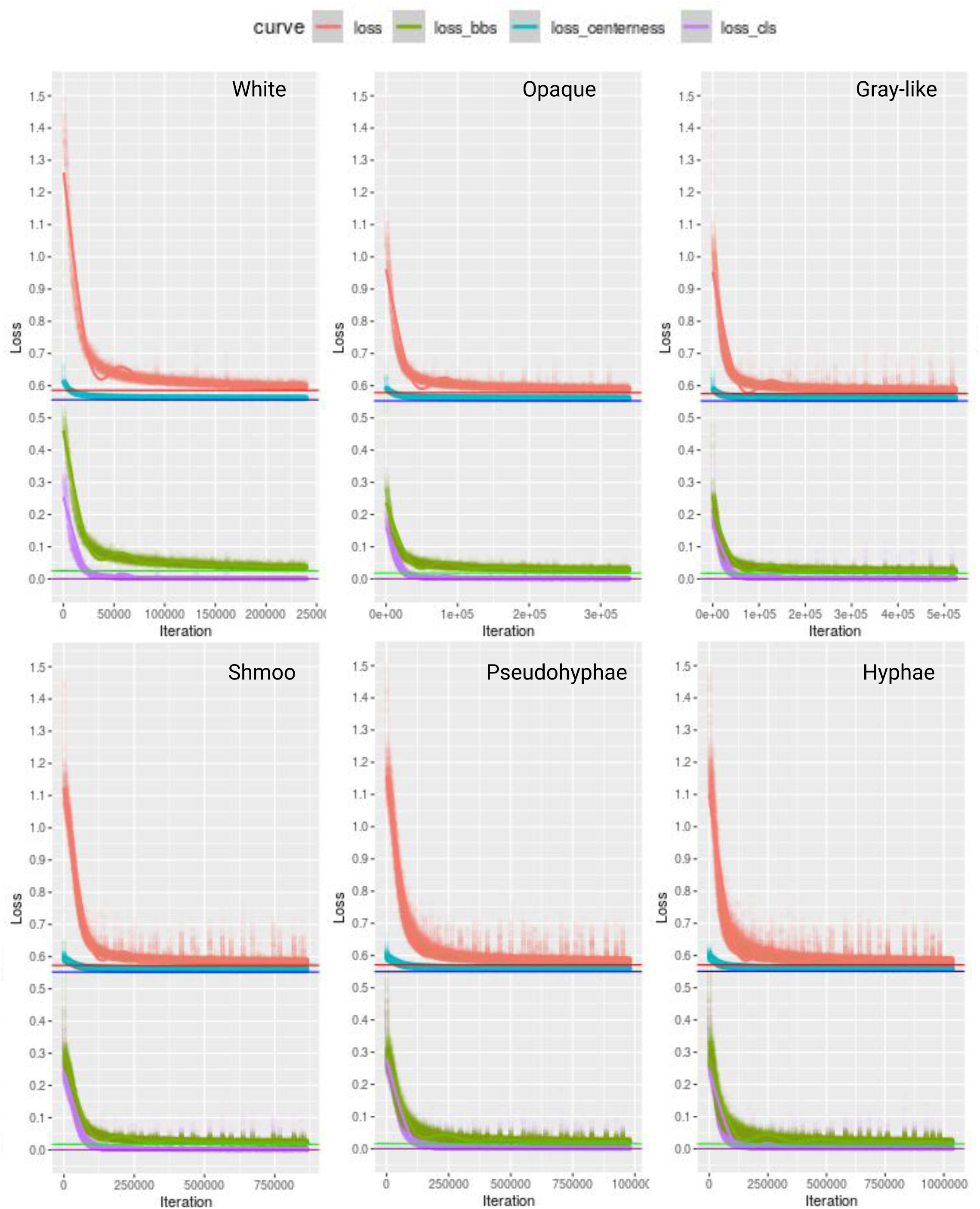
The curves for the three FCOS notions of loss across the six grades for the chosen Candescence classifier (τ=0.25) from Table 2. Although the number of iterations (x-axis) varies across the grades because of differences in the size of the learning set for each grade, in all cases this translates to a total of 5,000 epochs. From this, an epoch number of 1,000 was chosen, as it appears that convergence has been reached after one-fifth of the epochs. It is well-established that the center-ness loss converges to ∼0.57. All other losses are negligibly above 0.

**Supplemental Figure 3.**
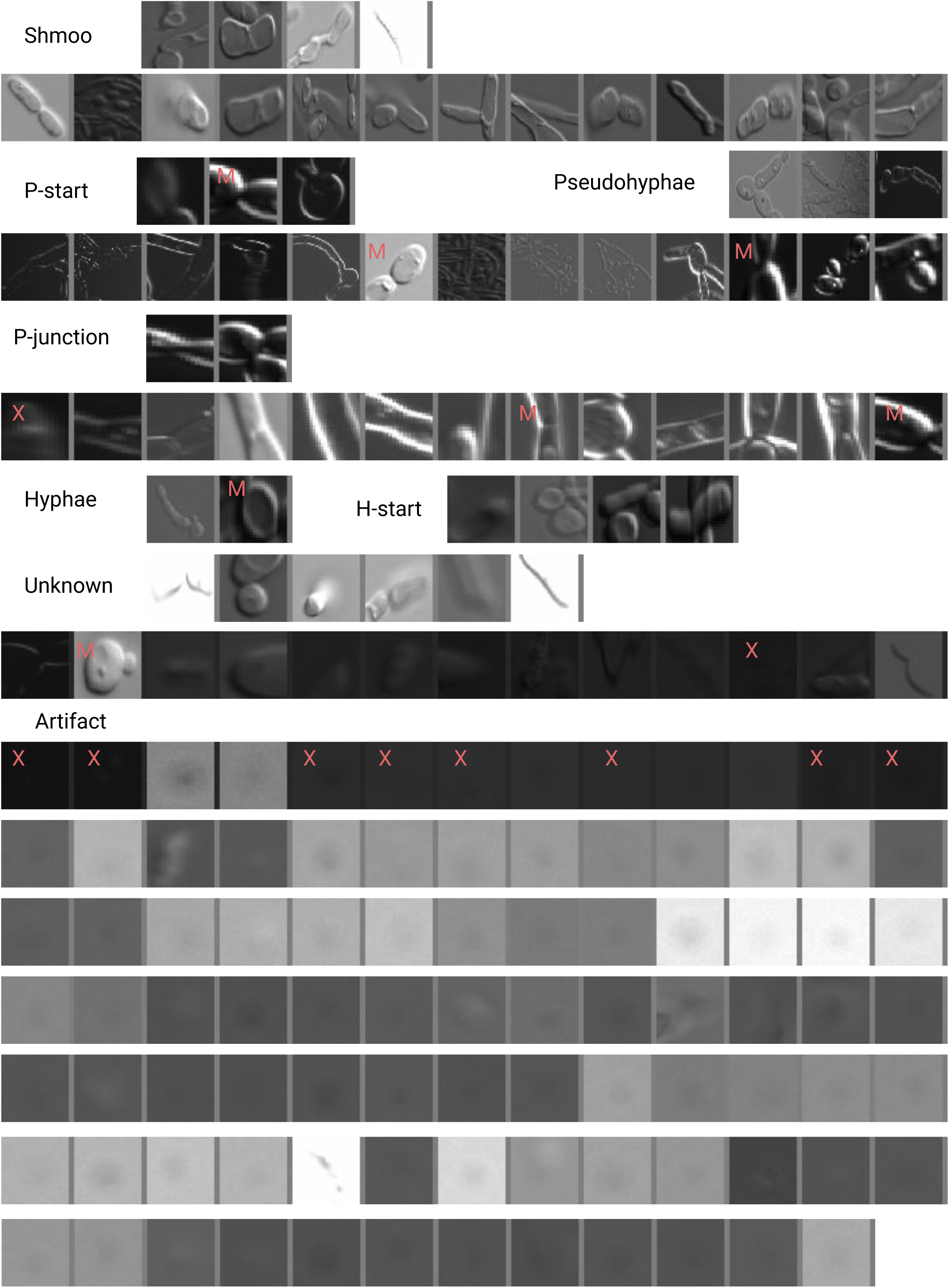
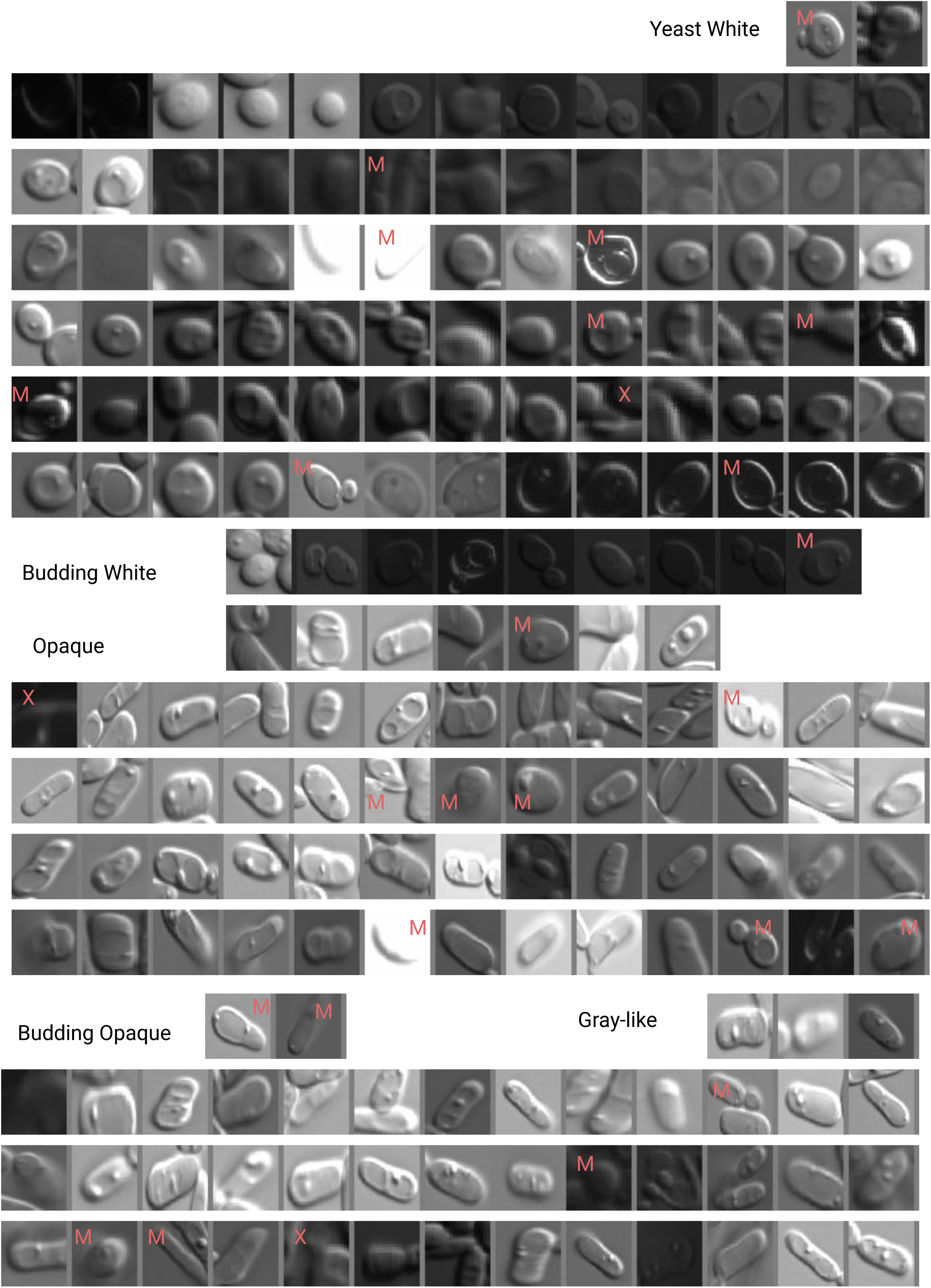
A compendium of all hallucinations (false positive object predictions) produced by Candescence. As in Figure 3(i), X indicates that we truly are not able to see an object (a true hallucination) and M indicates that Candescence was indeed correct to predict a bounding box at that location (missed by the human annotators) but its subsequent classification disagrees with our criteria. Subimages lacking an annotation indicate that Candescence correctly identified and classified the object, which represents a human error.

**Supplemental Figure 4.**
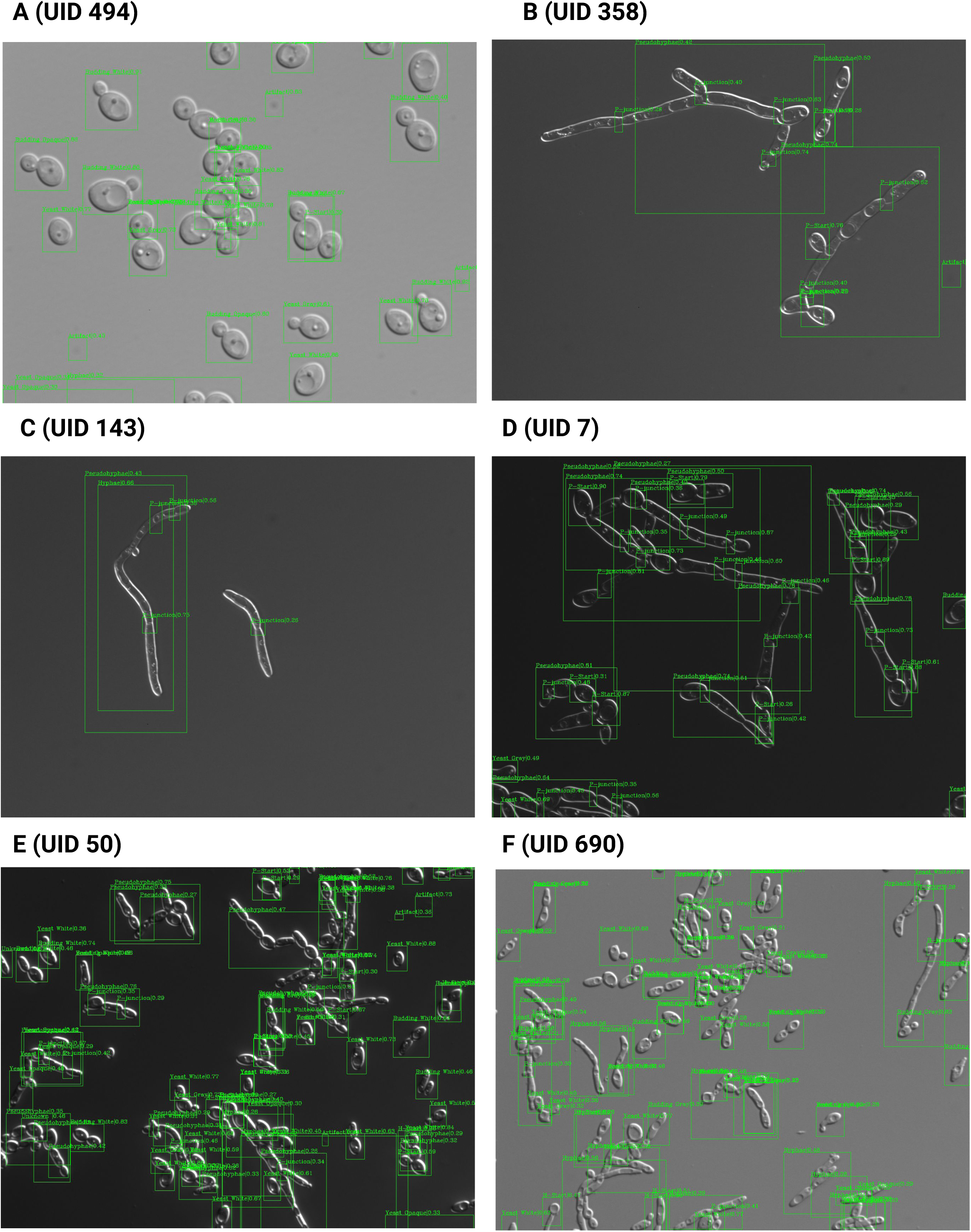
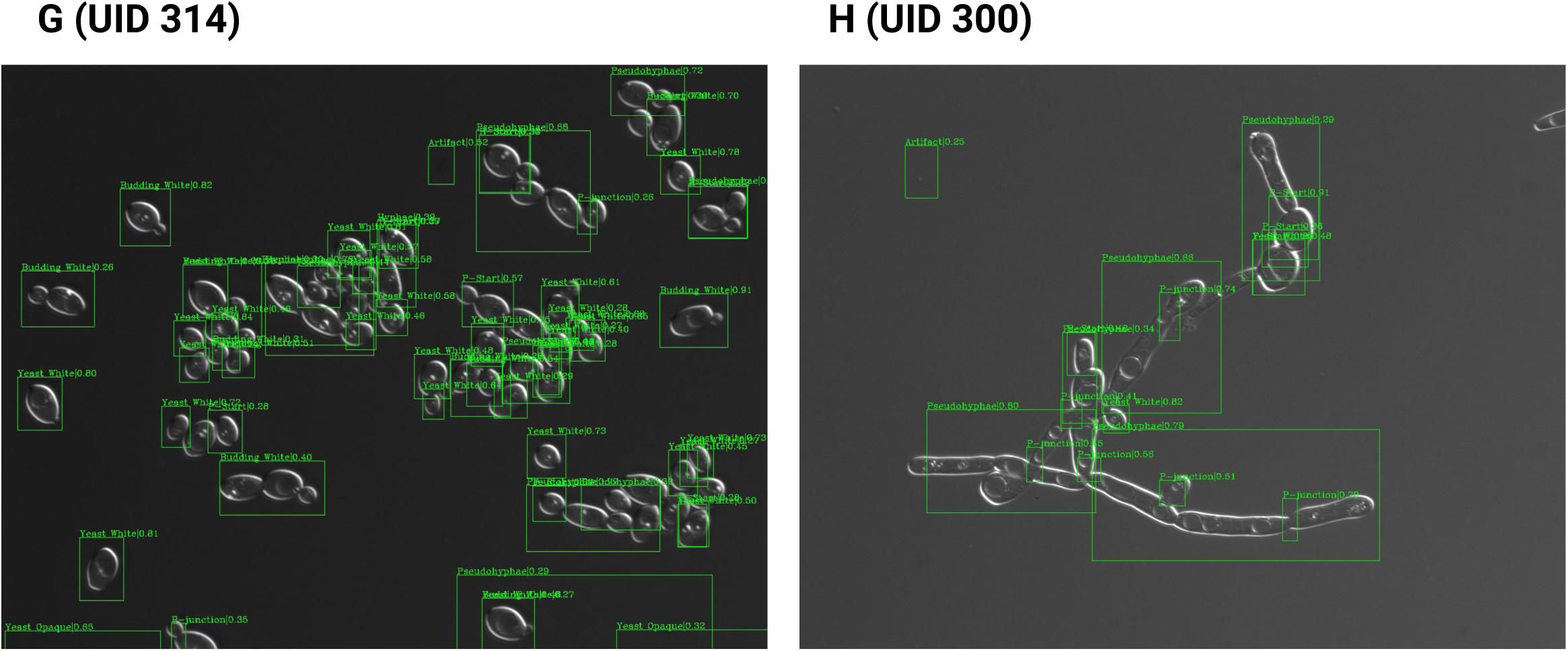
Exploration of the performance of Candescence on the test set. Each panel refers to one of ∼1,000 test set images with the UID available in Supplemental Table 1.

**Supplemental Figure 5.**
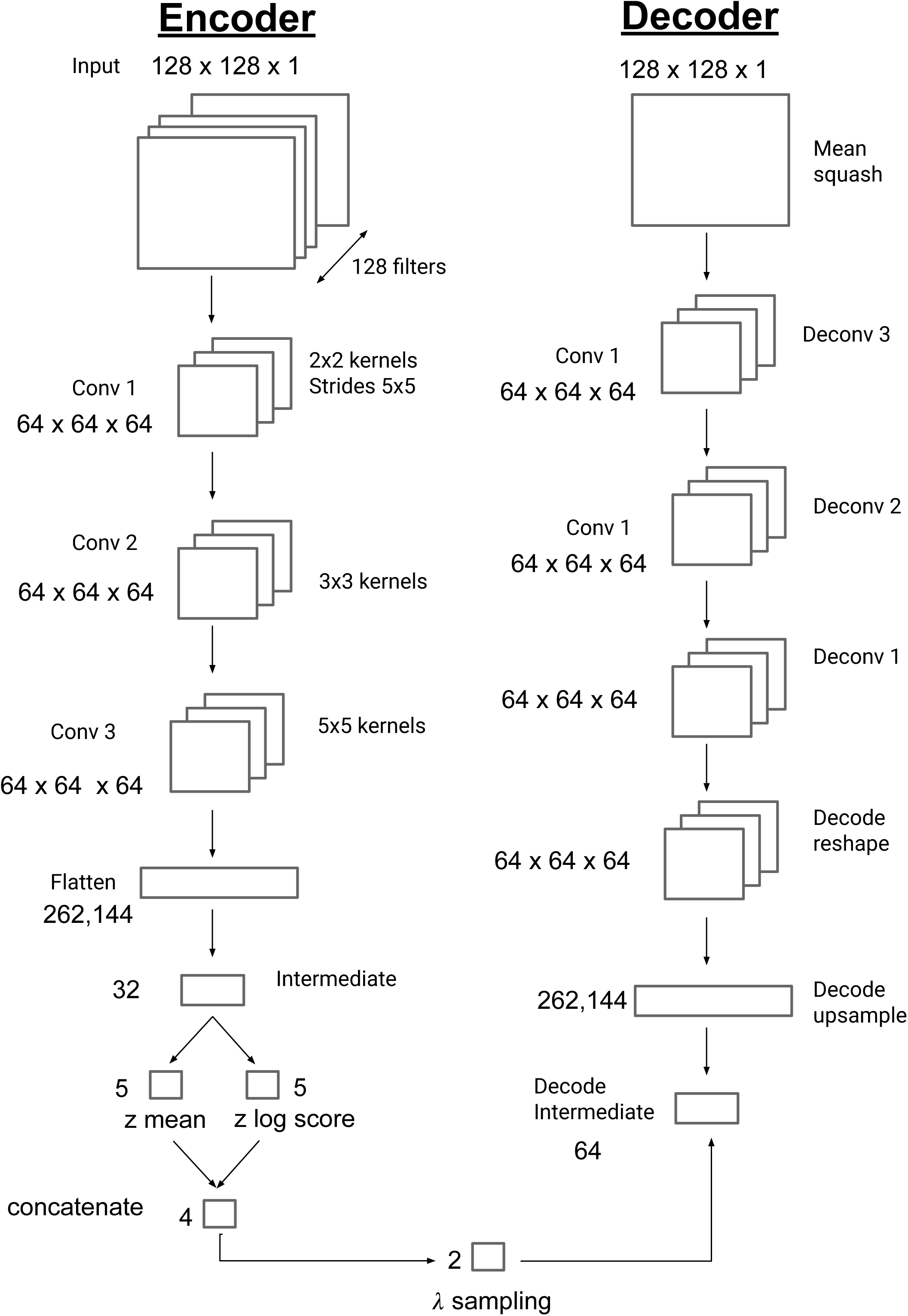
sketches the general design of our Keras R-based VAE. Input is a 128×128 subimage from the original images of the training and validation datasets. We opted here for a two-dimensional latent space for ease of visualization.

**Supplemental Figure 6.**
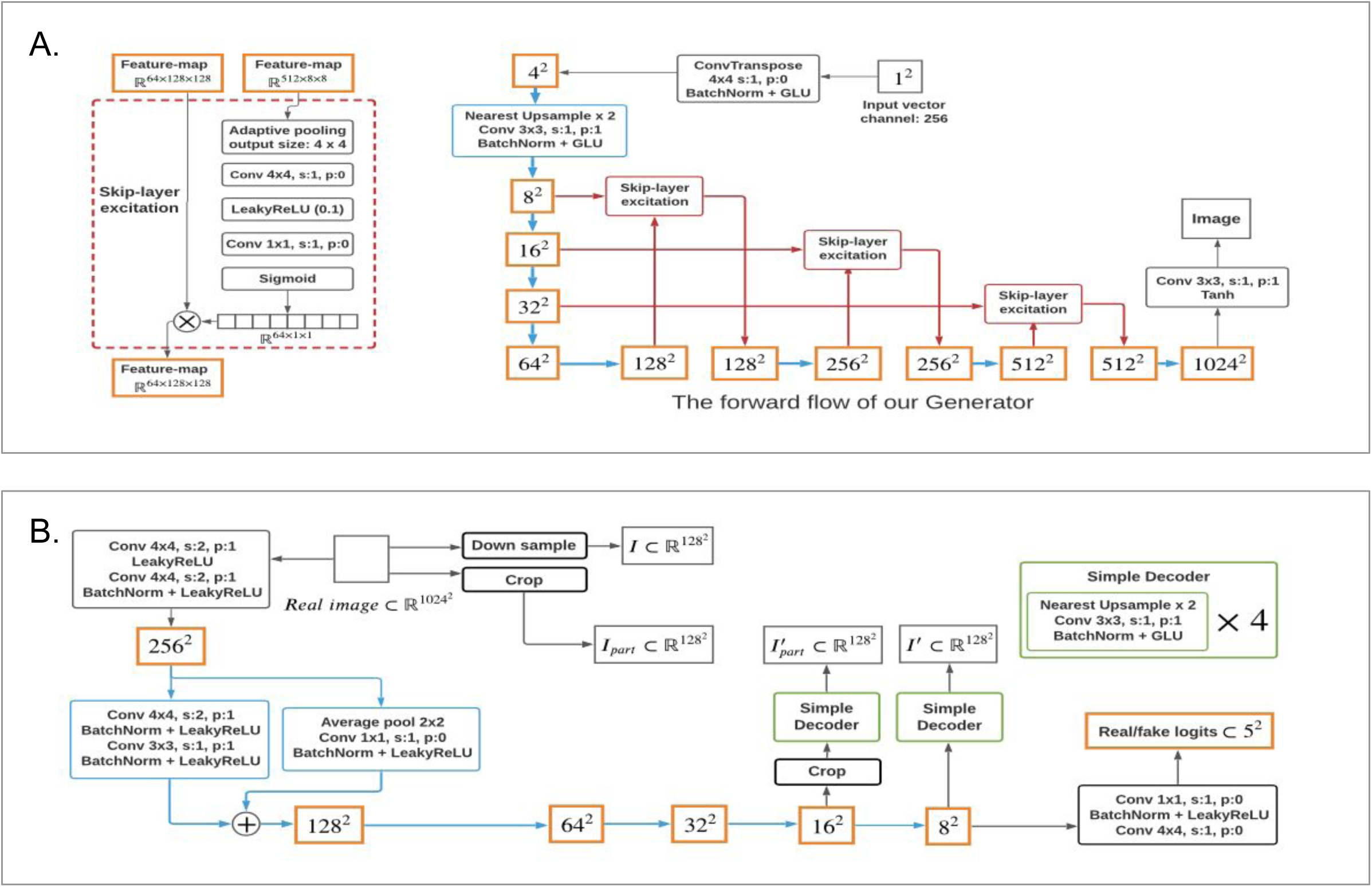
sketches the general design of our Generative Adversarial Network (GAN) trained on subimages as per the VAE. This design remains unchanged from the original presentation in Liu et al. (2021), although hyperparameters were fit using Varasana.

**Supplemental Figure 7.**
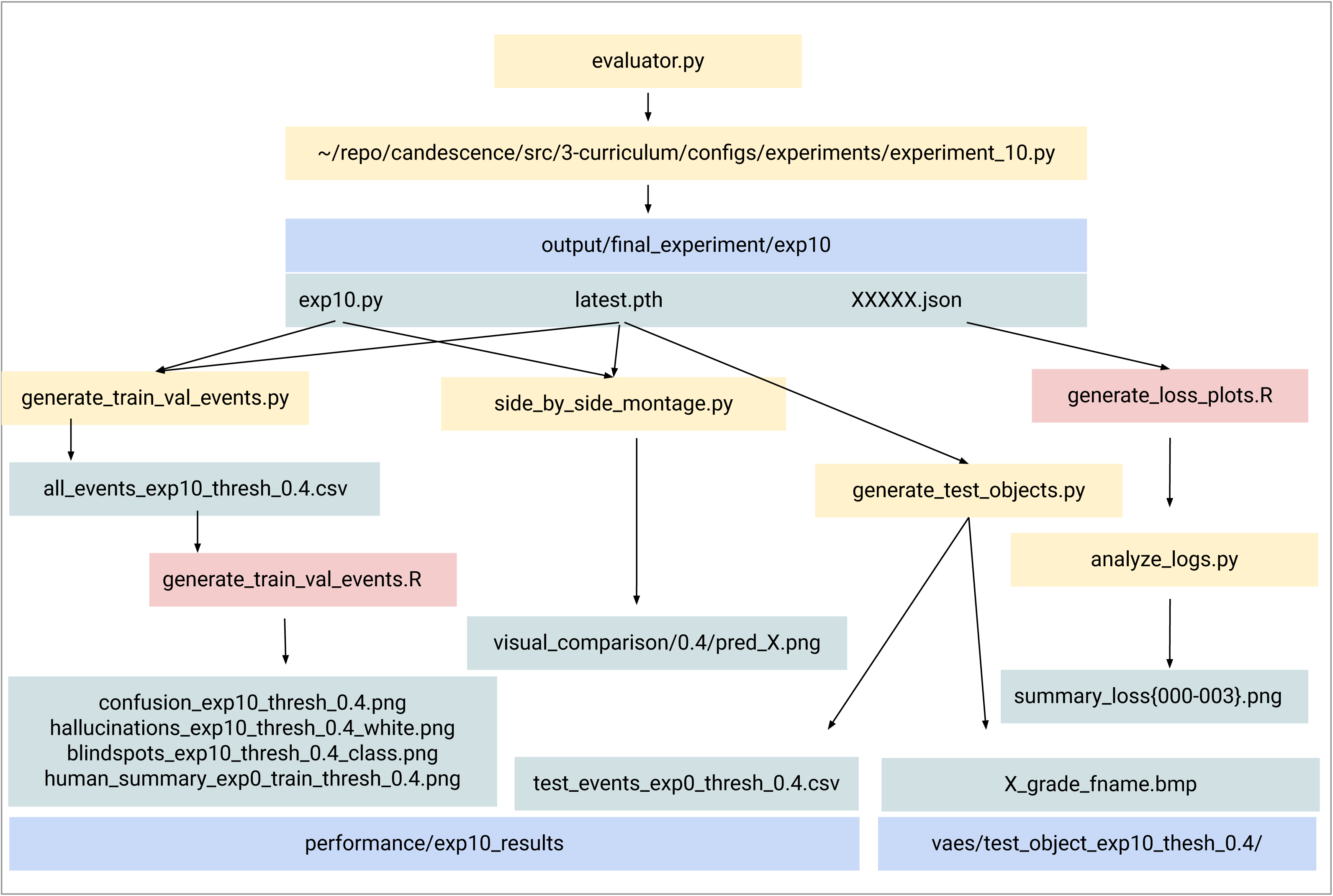
A description of the source code for (src/4-fcos-perf). The routines start with a classifier that has been learnt via experiment src/3-curriculum, and use this to measure its performance in different ways on the validation and test set. The scripts are written in Python or R, and make use of the FCOS implementation provided by MMDETECTION.

**Supplemental Table 1.** The Varasana learning set. UID is a distinct integer for each file. Type corresponds to a single colony that was photographed under the microscope and repetition refers to individual images taken of that colony. Gene target 1-3 describe the specific genetic modifications. ON under the Time column indicates overnight for 24 hours total. Columns S-AM provide the (manually assigned) number of cells per class per image across the training and validation datasets.

